# Disruption of recombination machinery alters the mutational landscape in plant organellar genomes

**DOI:** 10.1101/2024.06.03.597120

**Authors:** Gus Waneka, Amanda K. Broz, Forrest Wold-McGimsey, Yi Zou, Zhiqiang Wu, Daniel B. Sloan

## Abstract

Land plant organellar genomes have extremely low rates of point mutation yet also experience high rates of recombination and genome instability. Characterizing the molecular machinery responsible for these patterns is critical for understanding the evolution of these genomes. While much progress has been made towards understanding recombination activity in land plant organellar genomes, the relationship between recombination pathways and point mutation rates remains uncertain. The organellar targeted *mutS* homolog MSH1 has previously been shown to suppress point mutations as well as non-allelic recombination between short repeats in *Arabidopsis thaliana*. We therefore implemented high-fidelity Duplex Sequencing to test if other genes that function in recombination and maintenance of genome stability also affect point mutation rates. We found small to moderate increases in the frequency of single nucleotide variants (SNVs) and indels in mitochondrial and/or plastid genomes of *A. thaliana* mutant lines lacking *radA*, *recA1*, or *recA3*. In contrast, *osb2* and *why2* mutants did not exhibit an increase in point mutations compared to wild type (WT) controls. In addition, we analyzed the distribution of SNVs in previously generated Duplex Sequencing data from *A. thaliana* organellar genomes and found unexpected strand asymmetries and large effects of flanking nucleotides on mutation rates in WT plants and *msh1* mutants. Finally, using long-read Oxford Nanopore sequencing, we characterized structural variants in organellar genomes of the mutant lines and show that different short repeat sequences become recombinationally active in different mutant backgrounds. Together, these complementary sequencing approaches shed light on how recombination may impact the extraordinarily low point mutation rates in plant organellar genomes.

## INTRODUCTION

Nearly all eukaryotes rely on genes encoded in endosymbiotically derived mitochondrial genomes (mtDNAs) for cellular respiration. Plants and algae additionally rely on the endosymbiotically derived plastid genome (cpDNA) for photosynthesis. In several regards, land plant organellar genome evolution is atypical compared to mtDNA evolution in other eukaryotes (Smith and Keeling 2015). For one, plant organellar genomes have low nucleotide substitution rates relative to those in plant nuclear genomes and to those of many other eukaryotic mtDNAs. The low substitution rates of plant organellar genomes extend even to synonymous sites, which likely experience very little purifying selection, suggesting that the cause of the low evolutionary rates is a low underlying point mutation rate (Wolfe *et al*. 1987; Drouin *et al*. 2008).

Compared to the small mtDNAs typical in metazoans (generally below 20 kb) and in algae and fungi (with sizes ranging from approximately 13 to 96 kb and ∼20 to 235 kb, respectively), land plant mtDNAs are much larger with sequenced mtDNAs averaging 395 kb (Wu *et al*. 2022) and a known range extending from 70 kb to over 10 Mb (Boore 1999; Sloan *et al*. 2012; Skippington *et al*. 2017; Gualberto and Newton 2017; Sandor *et al*. 2018; Chen *et al*. 2019). Very little of this size variation stems from differences in coding capacity, as plant mtDNAs generally contain a subset of the same 41 protein-coding genes (Mower *et al*. 2012). Instead, the fluctuations in total mtDNA size primarily result from the acquisition and loss of noncoding DNA. Even closely related species possess very little shared noncoding sequence (Kubo and Newton 2008; Skippington *et al*. 2017). For example, a comparative analysis of the mtDNAs of two species within the Brassicaceae, *Arabidopsis thaliana* (367 kb) and *Brassica napus* (222 kb), revealed a mere 78 kb of shared sequence, most of which is coding (Handa 2003). Though size variation of cpDNAs is less extreme than in plant mtDNAs, variation still exists with 98.7% of sequenced land plant cpDNAs ranging from 100-200 kb in size (Xiao-Ming *et al*. 2017).

Plant organellar genomes also experience exceptionally high rates of structural mutation and rearrangement (Palmer and Herbon 1988). As a result, there is virtually no conservation of synteny between plant mtDNAs, as evidenced by the extensive rearrangements in alignments of mtDNAs from Col-0 and Ler ecotypes of *A. thaliana* (Stupar *et al*. 2001; Huang *et al*. 2005; Davila *et al*. 2011; Pucker *et al*. 2019; Zou *et al*. 2022). The structural instability in plant mtDNAs is partly explained by the presence of repeats of various lengths, which recombine frequently and give rise to multiple isomeric subgenomes with circular, linear and/or branched structures (Palmer and Herbon 1988; Alverson *et al*. 2011; Wynn and Christensen 2019). In fact, plant mtDNAs lack origins of replication, which help coordinate genome replication in many other eukaryotes, and are instead thought to replicate through break induced recombination (Gualberto and Newton 2017; Chevigny *et al*. 2020). Land plant cpDNAs are also recombinationally active but usually remain structurally conserved, albeit with some significant exceptions (Smith and Keeling 2015).

The seemingly disparate features of plant organellar evolution (i.e. high rates of recombination and low rates of point mutation) may be unified through a DNA repair mechanism reliant on recombination (Christensen 2014). This hypothesized mechanism hinges on the activity of the *mutS* homolog MSH1 (Abdelnoor *et al*. 2003), which is dual-targeted to mitochondria and plastids and has long been known to suppress non-allelic recombination between intermediate-sized repeats (50 to 600 bps) in the *A. thaliana* mtDNA (Martínez-Zapater *et al*. 1992; Arrieta-Montiel *et al*. 2009; Davila *et al*. 2011; Zou *et al*. 2022). Plant MSH1 is a chimeric fusion of a *mutS* gene with a GIY-YIG endonuclease domain (Abdelnoor *et al*. 2006) that has been proposed to introduce breaks in organellar DNA at the site of mismatches, which would then be repaired through homologous recombination (Christensen 2014, 2018; Ayala-García *et al*. 2018; Broz *et al*. 2022). Assays conducted on purified MSH1 *in vitro* have found that it has DNA binding and endonuclease activity with affinity for displacement loops (D-loops) (Peñafiel-Ayala *et al*. 2023).

We previously found support for a MSH1-mediated link between recombination and point mutations by using a high-fidelity Duplex Sequencing technique (Kennedy *et al*. 2014) to screen for single nucleotide variants (SNVs) and indels in *msh1* mutants (Wu *et al*. 2020). In that study, we also included a panel of mutants lacking functional copies of other genes involved in organellar DNA replication, recombination, and/or repair, including the recombination protein RECA3, the paralogous organellar DNA polymerases POLIA and POLIB, and the glycosylases UNG, FPG, and OGG (Wu *et al*. 2020). Compared to wild type (WT) lines, *msh1* mutants incurred SNVs at a ∼10-fold increase in mtDNA and a ∼100-fold increase in cpDNA, and increases in indel frequencies were even greater. In contrast, *recA3* mutants showed only a small (and marginally significant) increase in mtDNA mutation, and none of the other lines in the mutant panel showed a significant increase in SNVs or indels compared to WT plants (Wu *et al*. 2020).

Here, we investigate additional organellar genome repair proteins (WHY2, RADA, RECA1, OSB2) known to play a role in the suppression of non-allelic recombination in the *A. thaliana* organellar genomes. WHY2 is a mitochondrially targeted whirly protein that binds single-stranded DNA to inhibit recombination between small repeated sequences via micro-homology mediated end joining (MMEJ) (Cappadocia *et al*. 2010) and is also the most abundant protein in mitochondrial nucleoids (as measured in *A. thaliana* cell culture; Fuchs *et al*. 2020). RADA is a dual-targeted DNA helicase, which has been shown to accelerate the processing of recombination intermediates and promote mtDNA stability in *A. thaliana* (Chevigny *et al*. 2022). RECA1 is a plastid-targeted protein that has been proposed to act synergistically with plastid whirly proteins to promote plastid genome integrity either by facilitating polymerase lesion bypass or by reversing stalled replication forks (Rowan *et al*. 2010; Zampini *et al*. 2015). OSB2 is a plastid-targeted single-stranded DNA binding protein that has been shown to hamper microhomology-mediated end joining *in vitro* (García-Medel *et al*. 2021). Given that we previously saw a weak signal of increased mtDNA mutation in *recA3* mutants (Wu *et al*. 2020), we included another *recA3* mutant allele in this study. In addition to these newly generated mutant lines, we also present an extended analysis of Duplex Sequencing data from Wu *et al*. (2020) to understand how SNVs are distributed among genomic regions, strand (template vs. non-template) of genic regions, and trinucleotide contexts. Finally, we also performed long-read Oxford Nanopore sequencing on the mutant lines, allowing us to study structural mutations and rearrangements. Collectively, these analyses provide a detailed characterization of the effects of numerous recombination-related genes on point mutations and structural variants in plant organellar genomes.

## METHODS

### Generation and analysis of Duplex Sequencing libraries for SNV and indel detection

We obtained seeds for *A. thaliana osb2*, *radA*, *recA1*, *recA3*, and *why2* mutants from the Arabidopsis Biological Resource Center (Table S1). The generation of Duplex Sequencing data from mutants and matched WT controls (including crossing, plant growth, organelle isolation, DNA extraction, and library preparation) closely followed our previously described protocols (Wu *et al*. 2020). For each gene of interest, homozygous mutants were used as the paternal pollinators in crosses against WT maternal plants, which introduced ‘clean’ organellar genomes (i.e. never exposed to a mutant background) into the resulting heterozygous F1s. The presence of one WT allele in the F1 heterozygotes should be sufficient for WT-like organelle genome maintenance since the mutant alleles of the repair genes of interest are thought to act recessively (Shedge *et al*. 2007; Cappadocia *et al*. 2010; Rowan *et al*. 2010; Zampini *et al*. 2015; Wu *et al*. 2020; García-Medel *et al*. 2021; Chevigny *et al*. 2022). The heterozygous F1s were then allowed to self-cross and we identified three homozygous mutant and three homozygous WT F2s, which were also allowed to self-cross. Families of F3 seeds were grown together to obtain sufficient leaf tissue for organelle isolation and mutation detection via Duplex Sequencing.

The only notable differences between the methods in this study compared to Wu et al. 2020 were 1) we only isolated organelles for which the protein of interest is targeted (plastid: *OSB2*, *RADA,* and *RECA1*; mitochondrial: *RADA, RECA3, and WHY2*), whereas in Wu *et al.,* (2020) we isolated both organelles regardless of targeting. 2) We adjusted our Duplex Sequencing library construction protocol to obtain larger inserts by ultrasonicating the DNA for only 60 seconds (three bouts of 20 seconds, with 15 second pauses between each) and size selecting libraries with a 2% gel on a BluePippin (Sage Science), using a specified target range of 400-700 bp. 3) We implemented a new approach to filter spurious variant calls resulting from nuclear insertions of mtDNA and cpDNA (NUMTs and NUPTs) by comparing putative mutations directly against the *A. thaliana* nuclear genome (TAIR 10.2; Berardini *et al*. 2015) and the new assembly of the large NUMT on chromosome 2 (Fields *et al*. 2022), replacing the *k*-mer based NUMT/NUPT filtering approach described in Wu *et al*. (2020).

### Generation and analysis of nanopore sequencing libraries for structural variant detection

Nanopore libraries were produced from the same DNA samples that were used for Duplex Sequencing. Sequencing libraries were created following the protocol outlined in the Oxford Nanopore Technologies Rapid Barcoding Kit 96 (SQK-RBK110-96) manual (v110 Mar 24, 2021 revision) and were sequenced on MinION flow cells (FLO-MIN106) under the control of MinKNOW software v22.08.4 or 22.08.9. Multiplexed libraries from cpDNA samples were pooled and run on a single flow cell, whereas pooled mtDNA libraries were run on two flow cells. All runs were conducted for 72 hrs with a minimum read length of 200 bp. Data were processed using the Guppy Basecalling Software v6.3.4+cfaa134.

We sequenced three mutant replicates and one matched WT control for each gene of interest. Mutant lines for the cpDNA samples included *msh1* (CS3246)*, osb2, recA1,* and *radA* (only two *radA* mutants were sequenced due to a lack of DNA in mutant replicate 2), while mutant lines for the mtDNA samples included *msh1* (CS3246)*, recA3, why2,* and *radA*. The total sequencing yield (3.72 Gb) in our initial run of 15 cpDNA samples was an order of magnitude higher than our subsequent run with the 16 mtDNA samples (0.33 Gb). To increase mtDNA coverage we re-sequenced 12 of those mtDNA samples (all but the *msh1* mutants and matched WT control) in a third run, which had a similar low yield (0.42 Gb) to the second run. In all cases, samples were run on fresh flow cells as opposed to flow cells that had been washed for a second run. Because the *msh1* and *radA* mtDNA samples produced very little data (Table S4), we used the mtDNA contamination in the *msh1* and *radA* cpDNA samples in downstream analyses of the nanopore data.

To calculate mitochondrial and plastid read depth, we aligned the nanopore reads to the organellar genomes with minimap2 (version 2.24; Li 2018) and tabulated depth at each position with bedtools (version 2.30.0; Quinlan and Hall 2010). We calculated the average depth in 1000-bp sliding windows tiling the organellar genomes and plotted depth as a normalized mutant:WT ratio.

The nanopore reads were analyzed with HiFiSr (https://github.com/zouyinstein/hifisr), a software tool developed to identify structural variants using BLASTn alignments of long reads in plant organellar genomes (Zou *et al*. 2022). Because the tool was originally developed for PacBio HiFi reads, which are more accurate than nanopore reads, we required at least two independent nanopore reads to support putative indels. In addition, we constrained our analysis to reads with only one or two BLASTn hits, disregarding the reads with three or more BLASTn hits (which may originate from reads that span two or more recombined repeats). For reads with two BLASTn hits, we compared the breakpoints of putative recombination events with the repeats in the *A. thaliana* organellar genomes, which are reported in Tables S10 (mtDNA) and S28 (cpDNA) by Zou *et al*. (2022). We calculated recombination frequencies for each repeat pair as the number of recombined reads divided by the total number of repeat-spanning reads. To compute genome-wide repeat frequencies, we restricted the analyses to repeats that showed a total of at least ten mtDNA recombination reads across all replicates. Because cpDNA recombination events were much less common, we lowered the threshold to a minimum of three recombining reads per repeat for calculating recombinaiton frequencies. All of the matched WT controls were averaged for comparisons againsnt the mutant variant frequencies because we only sequenced one WT control for each gene of interest.

## RESULTS

### Duplex Sequencing coverage

We generated Duplex Sequencing libraries from DNA extracted from isolated organelles to test if genes involved in recombination-suppression also impact accumulation of SNVs and short indels in *A. thaliana* organellar genomes. Duplex Sequencing libraries were sequenced on a NovaSeq 6000 to produce between 30.6 to 139.1 million paired-end reads (2×150 nt) per library (Table S2). Processing the Duplex Sequencing libraries to collapse Illumina reads into consensus sequences and map them to organellar genomes resulted in coverage of 94.2 to 816.3× in the mitochondrial libraries (*radA, recA3,* and *why2*) and 234.2 to 1176.6× in the plastid libraries (*radA, recA1,* and *osb2*; Table S2).

### Increased SNV and indel frequency in *radA*, *recA1*, and *recA3* mutants

We compared variant frequencies of each mutant to the matched WT controls (two-tailed *t*-test) and found significant increases in SNV and indel frequencies in the *radA* mutants (p-values reported in Fig. 1). We also observed significant indel and weakly significant SNV increases in the *recA3* and *recA1* mutants in the mtDNA and cpDNA, respectively. We analyzed our previously generated *recA3* mutant from Wu *et al.,* (2020), which represents an independent mutant allele of *recA3*, and similarly found significant indel and weakly significant SNV increases in mtDNA (Fig. S1). In total, we detected 204 SNVs and 123 indels in the newly generated Duplex Sequencing libraries (File S1). Dinucleotide mutations involve neighboring sites both experiencing a substitution at the same time and are increasingly being recognized as an important type of mutation (Kaplanis *et al*. 2019). We assessed whether these mutations increase in frequency in any of the analyzed mutant backgrounds but found no significant differences relative to WT controls (Wilcoxon signed rank test, p > 0.05, Fig. S2).

**Figure 1.**
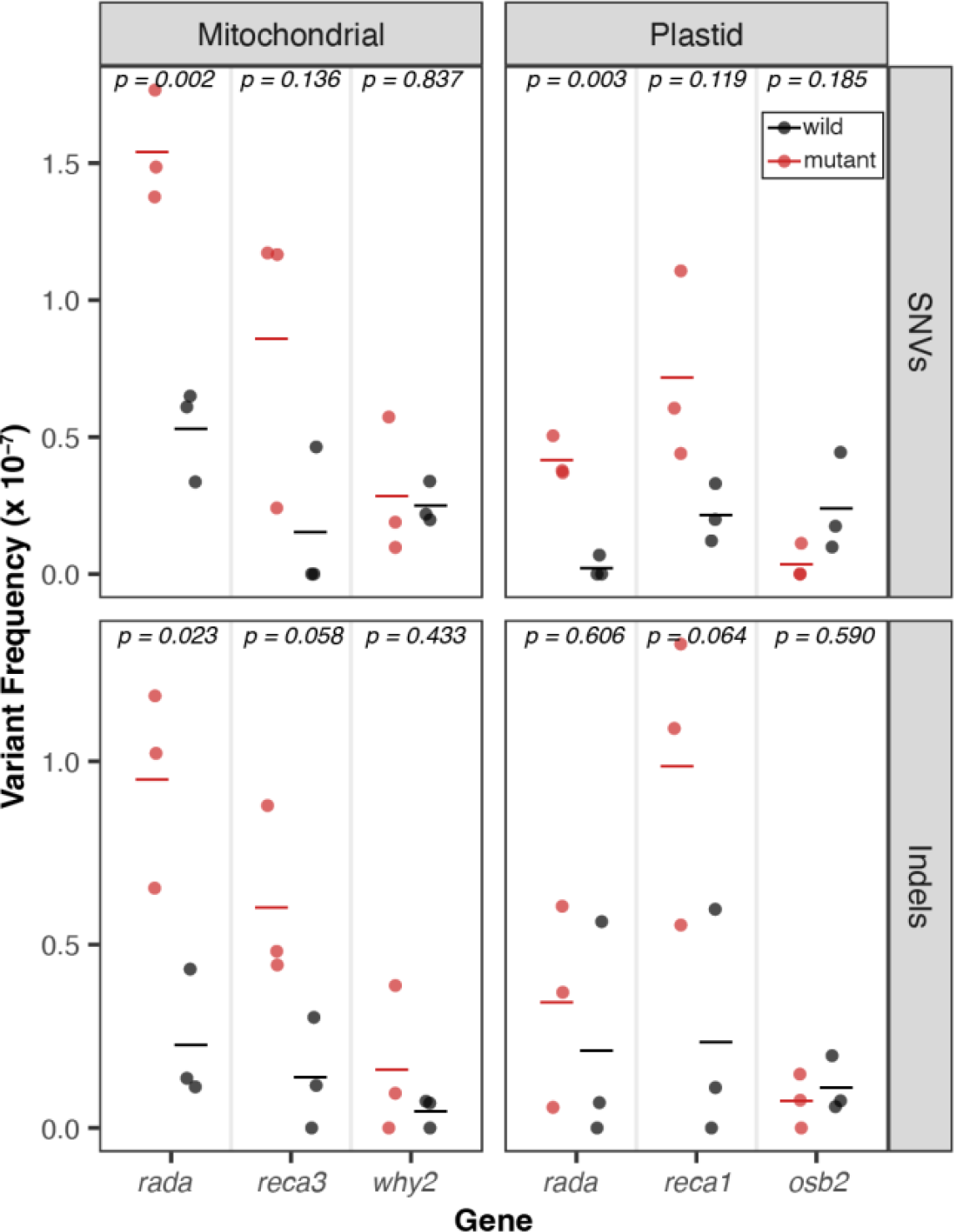
*De novo* point mutations measured with Duplex Sequencing. For each gene of interest (x-axis), mutant lines are plotted in red, and matched WT controls are plotted in black. The individual biological replicates are plotted as circles and group averages are plotted as dashes. Panels separate the data by genome; left column: Mitochondria and right column: Plastid, and by point mutation type; top row: SNVs and bottom row: indels. Variant frequencies (y-axis) were calculated as the total number of SNVs/total Duplex Sequencing coverage. P-values show the result of a two-tailed *t*-test comparing WT vs mutant mutation frequencies for each gene of interest.

### Decreased frequency of CG**→**TA transitions in the mtDNA of newly generated WT lines

The mutant lines assayed in both this study and in Wu *et al*. (2020) were sequenced with matched WT controls. Surprisingly, pooled WT SNV frequencies generated in the current study were lower than the pooled WT SNV frequencies from the Wu *et al*. (2020) dataset (2.8×10^-8^ vs. 1.7×10^-7^, *t*-test, p = 8.9×10^-12^), driven by a decrease in CG→TA transitions (*t*-test, p = 2.2×10^-10^; Fig. 2, File S1). To understand if the decreased SNV rate in the newly generated WT libraries (Fig. 2) resulted from the changes we made to our library preparation protocol, we created a Duplex Sequencing library following our new protocol using one of the original WT DNA samples from Wu *et al.,* (2020). This new library had an SNV rate of 1.57×10^-7^ which is in line with the SNV rates observed in the WT libraries from the 2020 study (Fig 2). In fact, the new SNV rate for this DNA sample was slightly higher than that of the original library (1.39×10^-7^). Given that the newly created libraries were all size selected on a BluePippin, which involves mixing the libraries with fluorescein labeled DNA as an internal standard for gauging DNA migration speed, we re-sequenced two stored libraries from Wu *et al.,* (2020) with and without size selection on the BluePippin. The inclusion of the sample without size selection on the BluePippin served as a control for the sample processed on the BluePippin and also as an independent test to understand if changes in the sequencing platform could be responsible (all samples were sequenced on a NovaSeq 6000, but the chemistry of the flow cells has been updated). These re-sequenced libraries had SNV rates typical of the old WT libraries of 1.97×10^-7^ (size selected library) and 1.47×10^-7^ (not size selected). Again, these values were slightly higher than the SNV rates from the original round of sequencing (1. 36×10^-7^ and 1.39×10^-7^, respectively).

**Figure 2.**
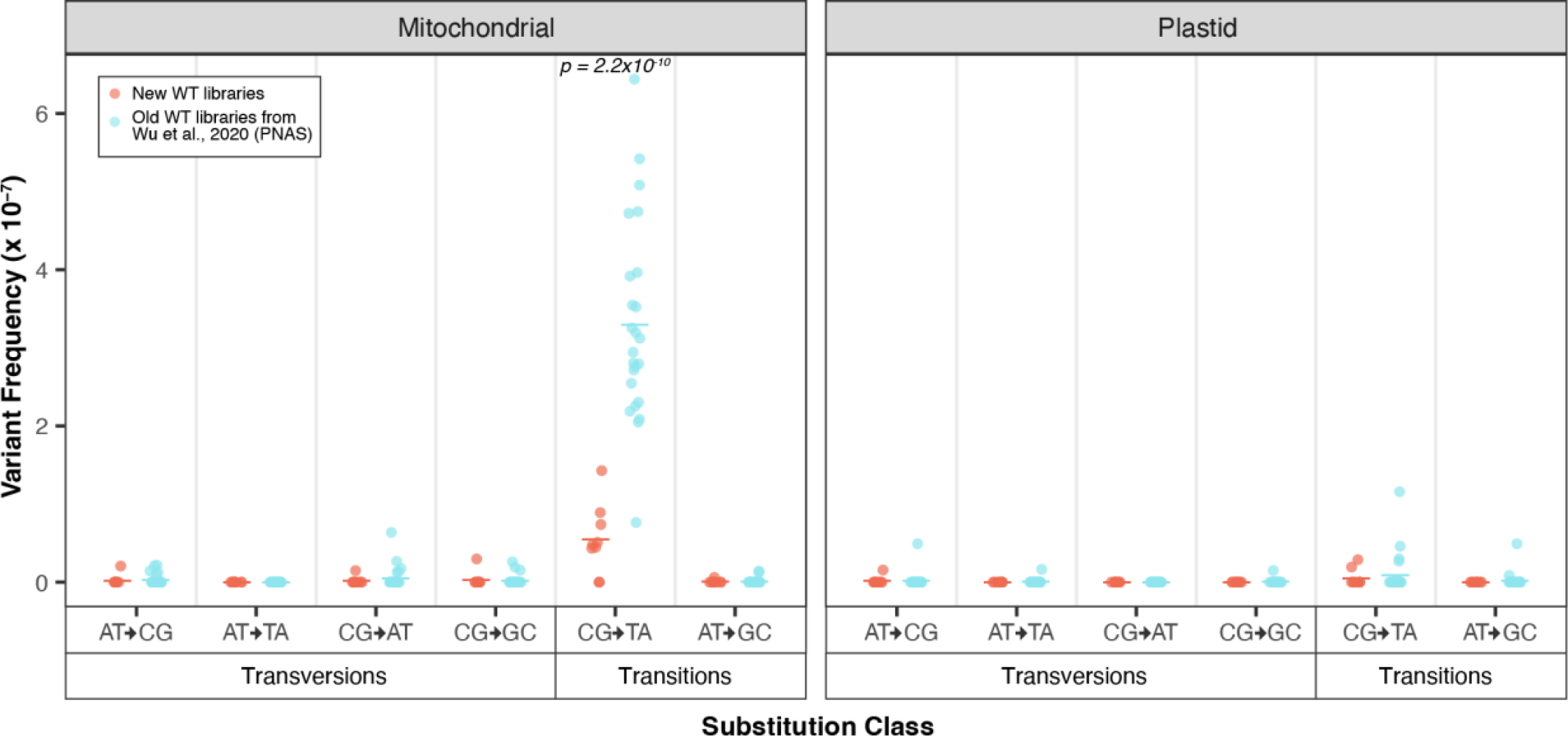
Comparison of the mutational spectrum of pooled WT controls from the current study (orange) vs. the WT controls from Wu *et al*. 2020 (blue). The two panels show the mitochondrial and plastid data and the x-axis separates substitutions type by transversions vs. transitions and further by the six types of substitutions. Individual biological replicates are plotted as circles while group averages are plotted as dashes. Only CG→TA transitions showed a significant increase in the old data set (two-tailed t-test; p=2.2×10^10^).

Therefore, it seems highly unlikely that the decreased SNV rate in the new WT libraries is associated with the changes we made to our library preparation protocol. Instead, these appear to be genuine differences in the DNA samples, perhaps due to unknown variation in the growth conditions or DNA extraction procedures between the two batches.

### SNV frequencies are similar among different genomic regions

To gain a deeper understanding of mutational process in the organellar genomes, we next turned our attention to the distribution of SNVs, focusing primarily on the *msh1* mutants and the pooled WT libraries from the Wu *et al*. (2020) study, given the larger number of mutations in those datasets. First, we assessed if the SNVs in *msh1* mutants and pooled WT libraries from Wu *et al*. (2020) are evenly distributed between intergenic, protein-coding (CDS), intronic, rRNA, and tRNA regions (Fig. 3) and found no significant differences among genomic regions (Kruskal-Wallis test, p > 0.05, Table S3) except in the WT plastid comparison, which is likely not biologically meaningful, given the small number of observed WT plastid SNVs (Fig. 2). Given that the vast majority of mtDNA SNVs in the Wu *et al*. (2020) WT dataset are CG→TA transitions, we separately tested if this class of substitutions is evenly distributed across regions and found significant differences (Kruskal-Wallis test, p = 0.0295), driven by a decrease in tRNA genes compared to intergenic sequences (pairwise comparisons with Wilcoxon rank sum test, p=0.0013). However, tRNA genes make up a small fraction of the genome and, thus, are subject to higher sampling variance, precluding any confident conclusions about whether they actually accumulate fewer CG→TA transitions than intergenic sequence.

**Figure 3.**
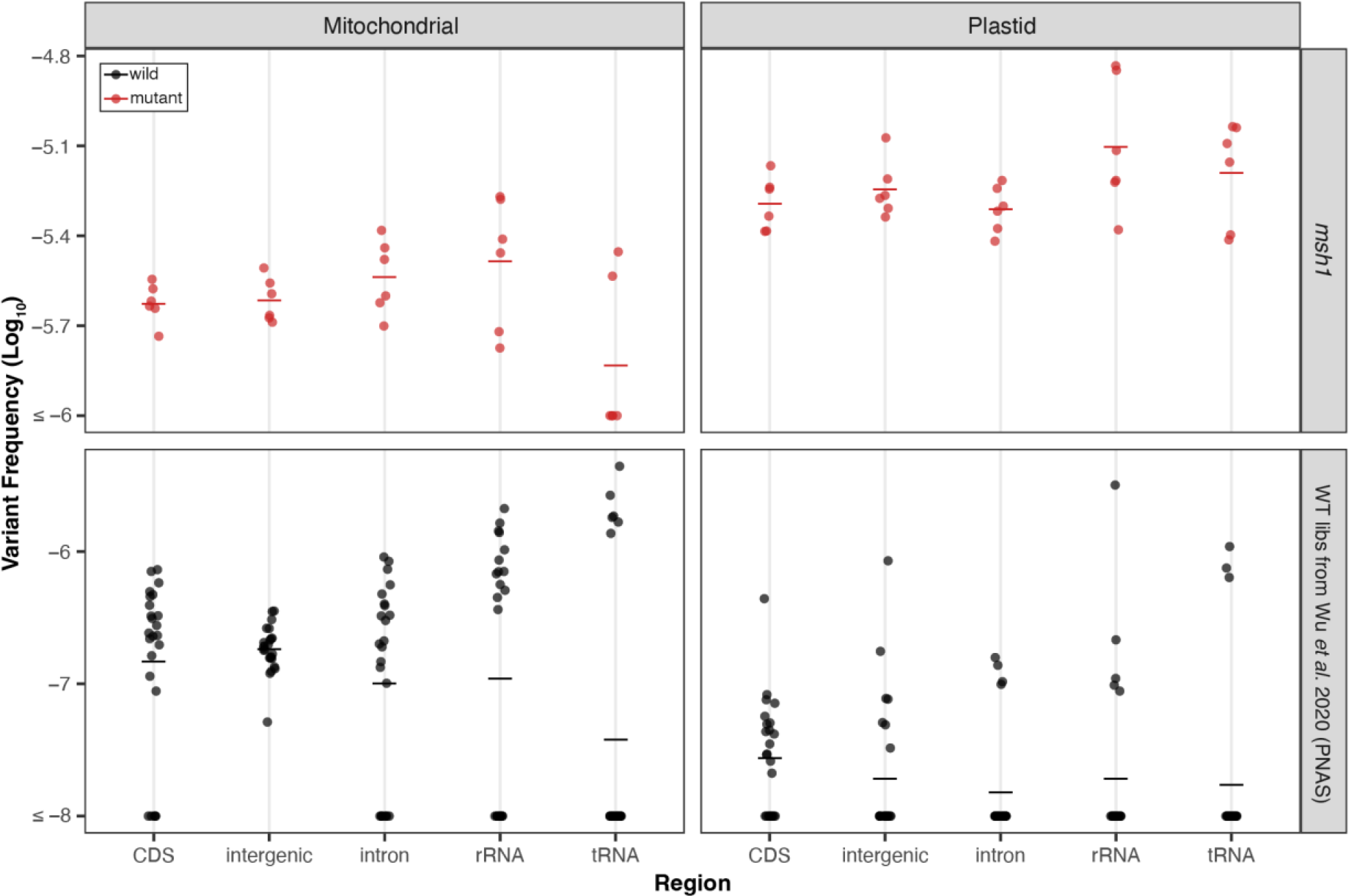
Distribution of WT (black) and *msh1* (red) SNVs (from Wu et al., 2020) across genomic region. The individual biological replicates are plotted as circles and group averages are plotted as dashes. Panels separate the data by genome; left column: Mitochondria and right column: Plastid, and by genotype with *msh1* mutants on top and WT on the bottom. Note the difference in y-axis scale for *msh1* mutants and WT. For each of the four panels, we performed a Kruskal-Wallis test and found no significant difference between genomic regions except the WT plastid panel (p = 0.022) where comparisons between regions are likely not biologically meaningful given the low number of WT plastid mutations. Note that for this and subsequent analyses of the *msh1* Duplex Sequencing data, we pooled the two null *msh1* alleles to increase statistical power.

### C**→**T substitutions are more common on the template strand in genic regions

Next, we performed a strand asymmetry analysis to understand if the SNVs in these datasets are evenly distributed on template vs. non-template (i.e., sense or coding) strands in the CDS, intronic, rRNA, and tRNA regions of the organellar genomes. The analysis of the CG→TA transitions from the Wu *et al*. (2020) WT dataset revealed that G→A substitutions are significantly enriched on the non-template strand of the DNA (paired Wilcoxon signed-rank test; p < 0.05 for CDS, rRNA and tRNA genes). Conversely, C→T substitutions predominately occur on the template strand, which is read by RNA polymerases during transcription (Fig. 4). This asymmetry is most striking in rRNA and tRNA genes, where every C→T substitution occurred on the template strand (25 in rRNA and 7 in tRNA). CG→TA transitions were also asymmetrically distributed between strands in genic regions of the Wu *et al*. (2020) *msh1* mutants (Fig. 5), though only in certain regions of the mtDNA (Fig. 5 top right panel), and not in the cpDNA (Fig. 5 bottom right panel). We also investigated strand asymmetries in the AT→GC transitions of the Wu *et al*. (2020) *msh1* mutants and found a trend toward more C→T substitutions on the template strand of plastid genes (Fig. 5 left panels). We did not investigate strand asymmetries for the other substitution classes in WT or *msh1* mutants because the small number of data points precludes meaningful comparisons between strands (see Fig. 5 of Wu *et al*. 2020).

**Figure 4.**
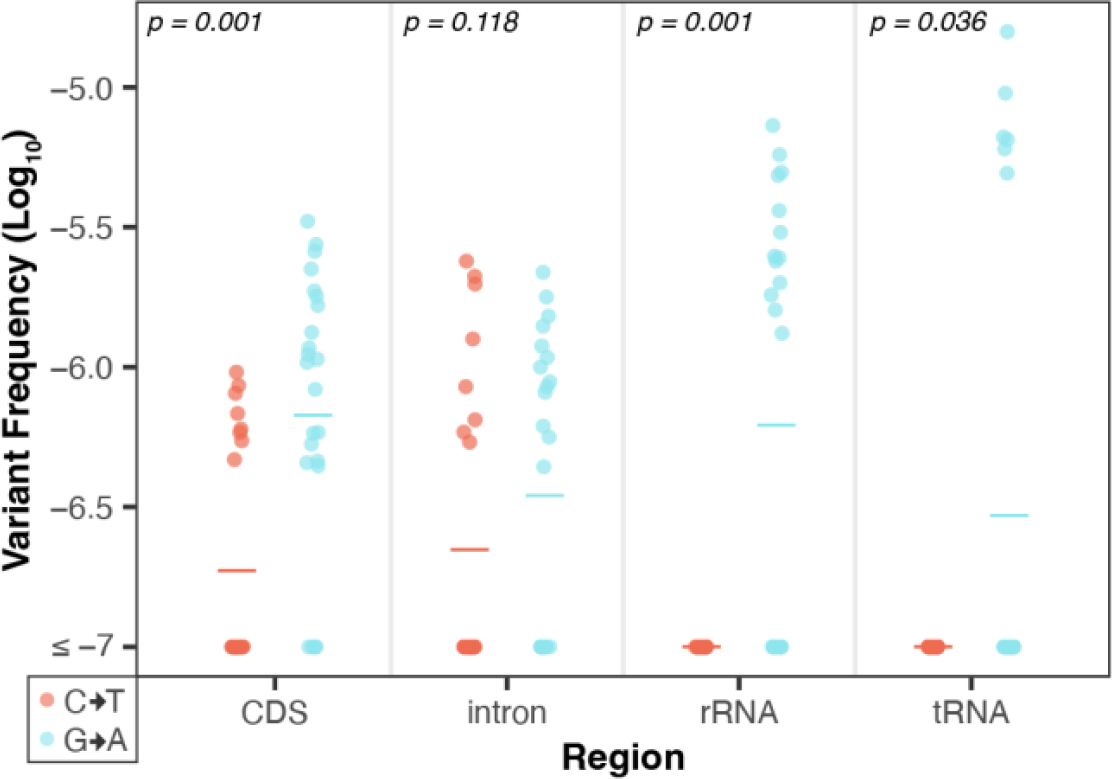
Strand asymmetry analysis of CG→TA transitions in the WT mtDNA Duplex Sequencing data from Wu *et al*. (2020). Shown are the log-transformed SNV frequencies (y- axis) of C→T (red) vs. G→A (blue) mutations on the non-template strand of all genes, separated by genomic region (x-axis). The individual biological replicates are plotted as circles and group averages are plotted as dashes. P-values show the result of paired Wilcoxon tests comparing the complementary substitution classes in each genomic region. In all but intronic regions, G→A substitutions are significantly higher on the non- template strand (conversely, C→T substitutions are significantly higher on the template strand). Strikingly, in all of the observed CG>TA transitions in the rRNA and tRNA genes the C→T substitution occurred on the template strand (i.e., all the G→A substitutions occurred on the non-template stand) .

**Figure 5.**
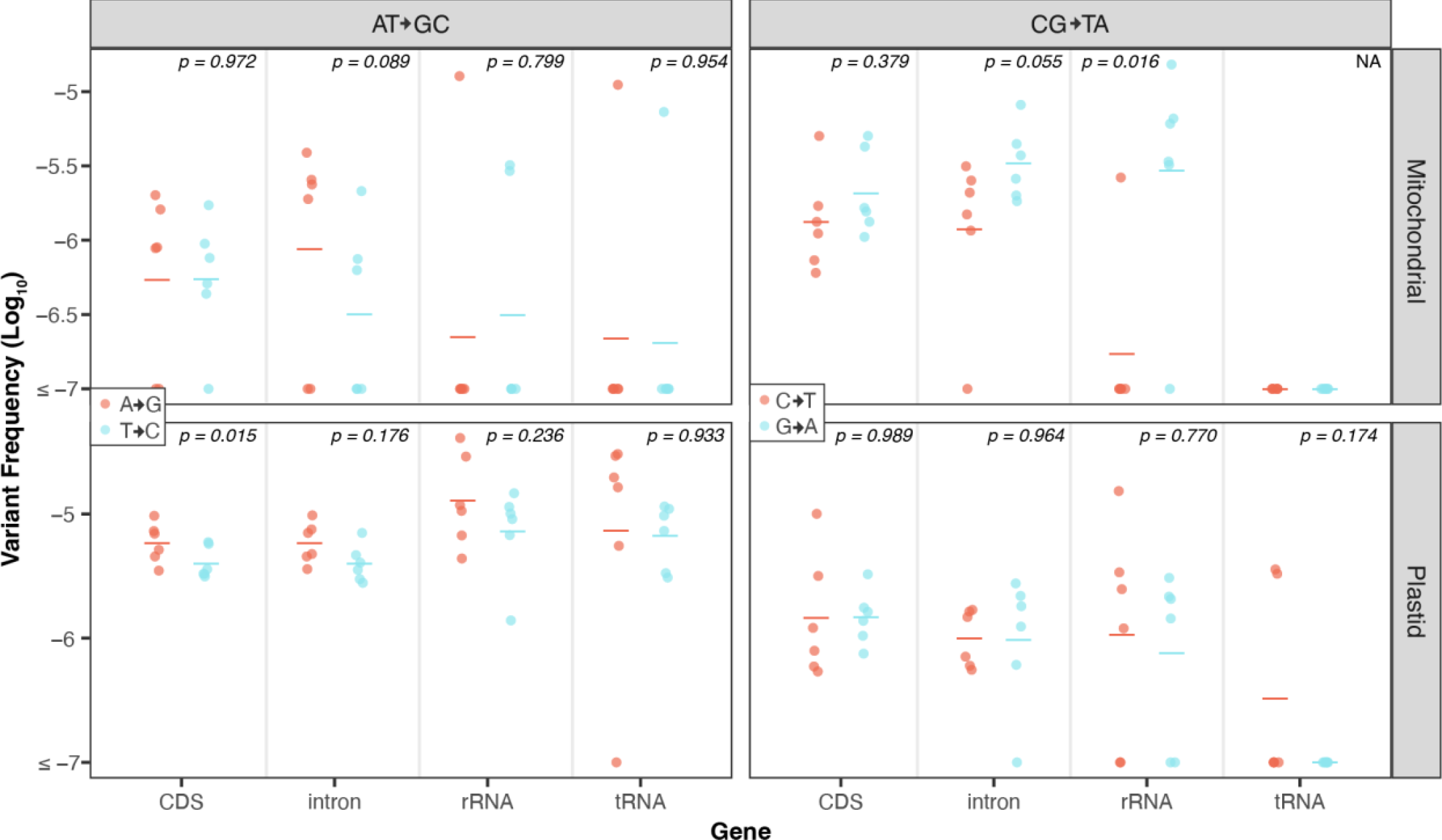
Strand asymmetry analysis of CG→TA and AT→GC transitions in the *msh1* Duplex Sequencing data from Wu *et al*. (2020). Shown are the log-transformed SNV frequencies (y- axis) of mutations on the non-template strands of all genes with complementary substitution types designated by color (see figure legends for colors of specific substitution types). The individual biological replicates are plotted as circles, and group averages are plotted as dashes. The panels divide the data by transition type, with AT→GC transitions on the left and CG→TA transitions shown on the right, and by genome, with mitochondrial data on the top and plastid data on the bottom. Transversions were not analyzed because there were relatively few observed mutations of this type in the *msh1* duplex data. P-values show the result of paired *t*-tests comparing the complementary substitution classes in each genomic region.

### CG**→**TA transition frequencies vary depending on trinucleotide context

To understand how surrounding nucleotides impact SNV accumulation in plant organellar genomes, we performed a trinucleotide analysis, again focusing on CG→TA transitions in WT and both transition types in *msh1* mutants, due to a lack of data in other substitution classes. In the WT dataset (Wu *et al*. 2020), we found that CG→TA transitions are 8.4-fold more common in the mtDNA and 3.7-fold more common in the cpDNA when the C is 3′ of a pyrimidine (Fig. 6). Interestingly, this same trinucleotide context (5′ pyrimidine) is not enriched for CG→TA transitions in the *msh1* mutant data. Instead CG→TA transitions are 3.0-fold more common when the C is 5′ of a G in the *msh1* mutants (Fig. 7 right panels).

**Figure 6.**
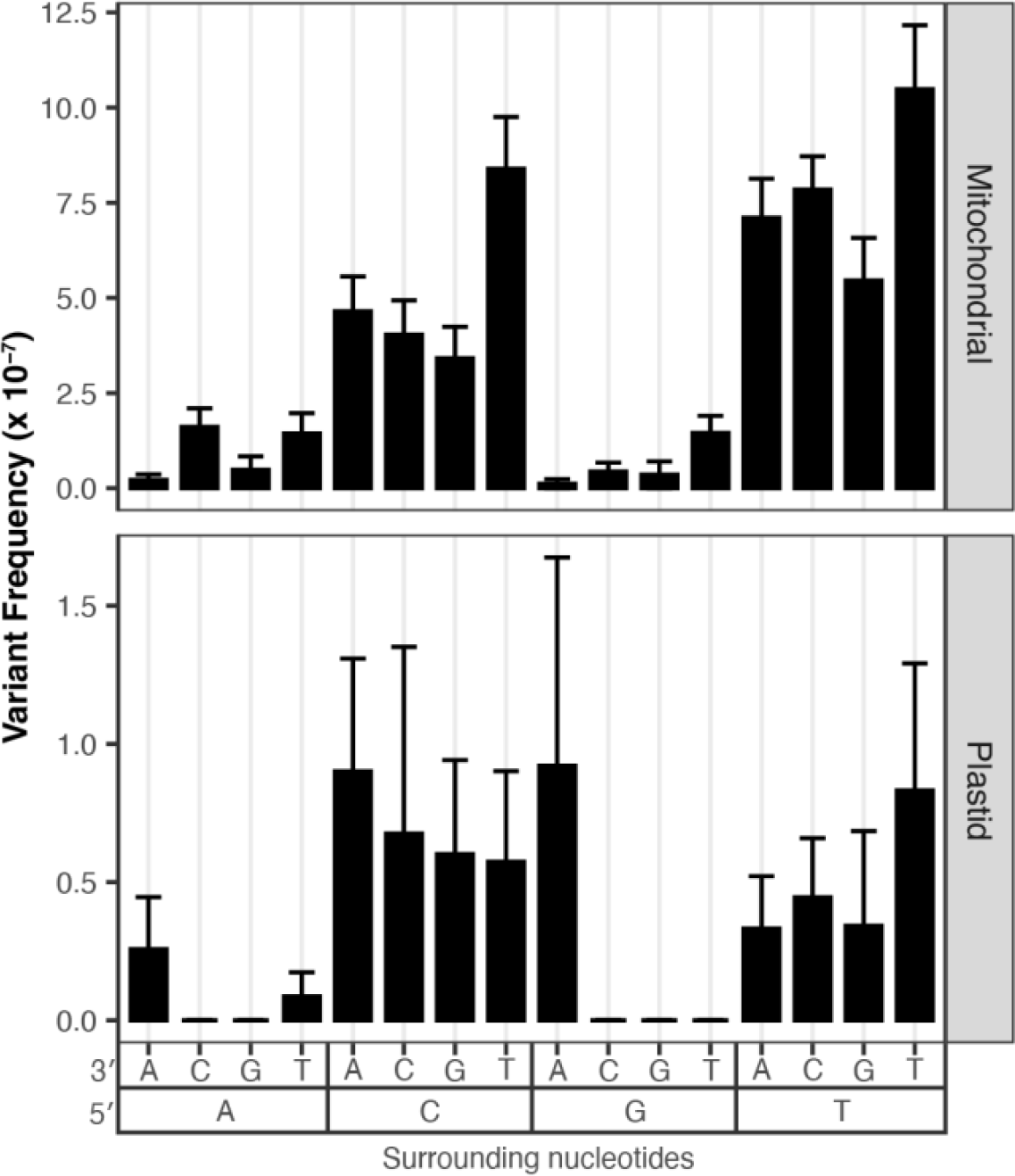
Analysis of surrounding nucleotides on C→T transition frequencies in the WT Duplex Sequencing data from Wu *et al*. (2020). The panels divide the data based on genome with mitochondrial data on the top and plastid data on the bottom, note the difference in the y-axis scale, as CG>TA were less frequent in the plastid. The x-axis captures the trinucleotide context with downstream nucleotides displayed next to the 3′ and upstream nucleotides display next to the 5′. The data suggest that trinucleotide contexts with upstream pyrimidines (5′ CCN 3′ and 5′ TCN 3′, where N is any nucleotide) have increased frequencies of C>T substitutions.

**Figure 7.**
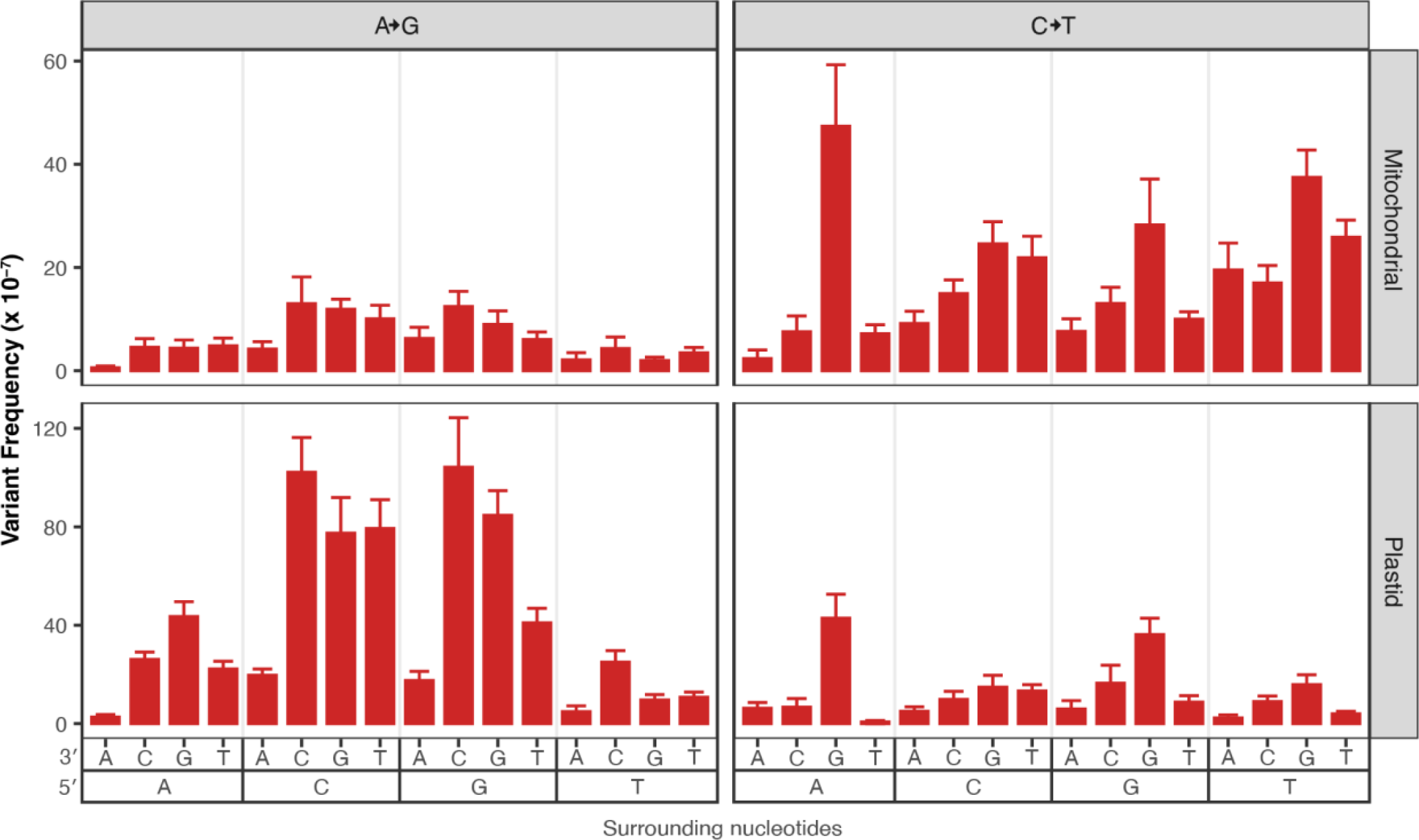
Analysis of surrounding nucleotides on A→G and C→T transition frequencies in the *msh1* Duplex Sequencing data from Wu *et al*. (2020). The panels divide the data based on substitution type (A→G substitutions on the left and C→T substitutions on the right) and by genome (mitochondrial data on the top and plastid data on the bottom). The x-axis captures the trinucleotide context with downstream nucleotides displayed next to the 3′ and upstream nucleotides display next to the 5′. The A→G data suggest that trinucleotide contexts with downstream Cs (5′ NAC 3′) have increased frequencies of A→G substitutions. The C→T data suggest that trinucleotide contexts with downstream Gs (5′ NCG 3′) have increased frequencies of C→T substitutions.

Meanwhile AT→GC transitions are 1.8-fold more common when the A is 5′ of a C (Fig. 7 left panels). In all cases, these trinucleotide mutation frequencies are normalized by the total coverage of a given trinucleotide context so that the values are not inflated in trinucleotides that are relatively common in the mtDNA.

### Chloroplast extractions produced an order of magnitude more nanopore sequencing data than mitochondrial extractions

We next generated long-read Oxford Nanopore libraries to gain a deeper understanding of how the genes in our panel impact plant organellar genome stability. Unexpectedly, the libraries produced from the mitochondrial isolations sequenced poorly compared to the plastid-derived libraries (see methods), so we investigated cross-organelle contamination (mtDNA molecules in the plastid-derived samples and cpDNA molecules in mitochondrially derived samples) to understand if poor mtDNA sequencing performance was inherit to the mtDNA or associated with differences in the organellar isolation methods. The level of mtDNA contamination in the plastid-derived nanopore libraries is similar to the level of contamination in the Duplex Sequencing libraries (Fig. S3). The average median read length of the mitochondrial derived nanopore libraries is about 2.5- fold higher than the average median read length of the plastid-derived libraries (2.48 kb vs. 1.08 kb, respectively). In the plastid derived nanopore libraries, the median lengths of the contaminating mtDNA reads tend to be slightly longer than the median lengths of native cpDNA reads (average median lengths of 1.17 kb vs 0.98 kb, respectively), though there is substantial variation between samples (Fig. S4). In the mitochondrially derived libraries, the contaminating cpDNA and native mtDNA median read lengths show more correlation (average median lengths of 2.41 kb and 2.56 kb, respectively; Fig. S4).

These analyses suggest that the difference in yields for the different nanopore runs is likely related to differences in the organellar isolation methods. One unique feature of the mitochondrial isolation protocol is the use of a DNase I treatment to remove contaminating nuclear and plastid DNA molecules (Wu *et al*. 2020). It is possible that this treatment results in nicking of the mtDNA that interrupts the molecules as they are threaded through the nanopore in a single-stranded fashion. Such nicking would not be expected to disrupt Duplex Sequencing library creation since the first step of making Duplex Sequencing libraries is to break DNA into small fragments via ultrasonication.

However, this explanation is somewhat inconsistent with the 2.5-fold greater median read length in the mitochondrially derived nanopore libraries. Fortunately, the contaminating mtDNA derived reads in the *msh1* and *radA* cpDNA sequenced samples provided sufficient mtDNA coverage for analyzing structural variation in the mtDNA (Table S4, Figure S3 left panel).

### Repeat-mediated recombination drives distinct patterns of mtDNA instability in *msh1*, *radA,* and *recA3 mutants*

Given the known role of recombination-related genes in maintaining organellar genome copy number and structural stability (Arrieta-Montiel *et al*. 2009; Davila *et al*. 2011; Miller-Messmer *et al*. 2012; Chevigny *et al*. 2022; Zou *et al*. 2022), we analyzed the ratio of mutant coverage to WT coverage to characterize structural perturbations on a genome-wide level (Fig. 8). We see distinct variation patterns in the mtDNA coverage in *msh1, radA* and *recA3* mutants, consistent with the expected structural effects of these genes (Fig. 8) and similar to previously documented coverage patterns (Wu *et al*. 2020; Chevigny *et al*. 2022). In contrast, the *why2* coverage does not deviate from WT coverage, suggesting there is no substantial and consistent structural effect of losing *why2.* In *recA3*, the nanopore and Duplex Sequencing lines are tightly correlated, while the nanopore data tends to show greater variance in the *msh1, radA*, and *why2* plots, perhaps because of the lower nanopore coverage in those samples (Table S5; Figs. S6 and S67). Interestingly, *radA* and *recA3* share many major coverage peaks and valleys, suggesting genome structure is perturbed in similar ways in these mutants (Fig. 8, Figs S6 and S7). Compared to the mitochondrial samples, the cpDNA samples display much less coverage variation (Fig. S5), with a notable exception in the *recA1* nanopore data. However, inspection of the coverage in the individual cpDNA replicates (Fig. S8) reveals depth irregularities in the WT control compared to the other WT samples. Regardless, the *recA1* Duplex Sequencing data does not show any depth variation along the cpDNA, so the nanopore result does not appear to reflect a biological effect on cpDNA structure. One other intriguing pattern in the cpDNA plots is an apparent correlation in peaks and valleys in *radA* and *osb2* in the Duplex Sequencing data (most notable is the shared valley at 112 kb). However, inspection of the individual *recA1* mutant and matched WT control replicates (Fig. S9) reveals all samples have a dip at 112 kb and the dip is more pronounced in one or more of the *osb2* and *radA* mutants. Given the large number of PCR cycles used to amplify the Duplex Sequencing libraries (19 cycles) the unified movement of all replicates is likely explained in part by amplification bias in AT or GC rich regions. Therefore, variation in amplification bias may result in lower coverage of AT or GC rich regions, so these patterns are likely not biological.

**Figure 8.**
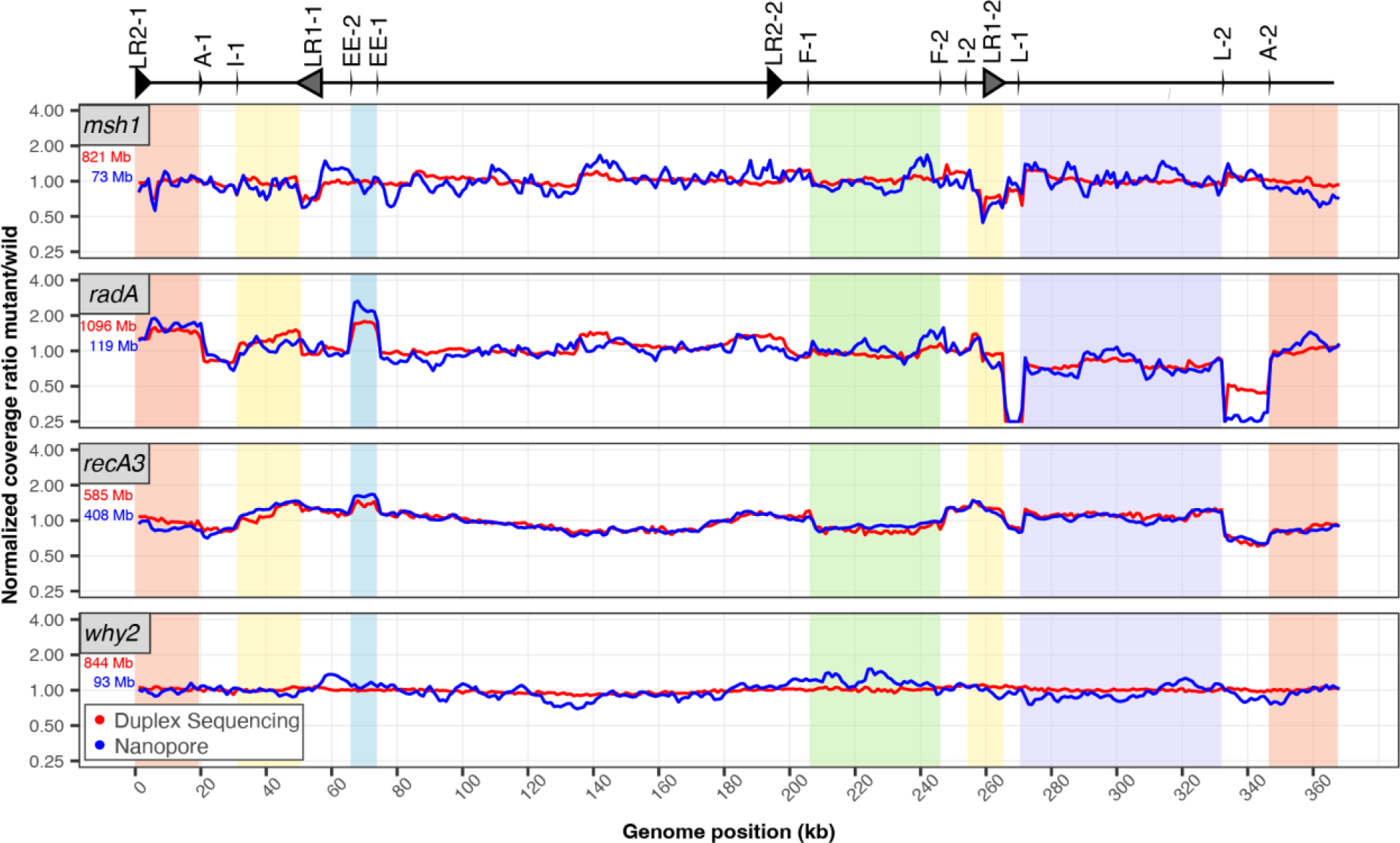
Normalized coverage of mitochondria genomes in mutant lines of interest . Coverage of each Duplex Sequencing (red) or nanopore (blue) library was calculated in 1000-bp windows. Mutant coverage was pooled and divided by WT coverage and the resulting ratios were normalized to 1 for plotting. The total amount of sequencing data used to generate each plot is shown in the top left corner of each panel (red=Duplex Sequencing and blue=nanopore) and is included to highlight the instances where disagreement between the Duplex Sequencing and nanopore lines may be explained by increased variance in the nanopore sample due to lower mtDNA coverage. Repeats that are likely important for driving coverage variation across the mtDNA are plotted above (also see, Table 1) according to Figure 6 of Chevigny *et al*. 2022. Regions with altered stoichiometry and flanked by repeats are shown as colored blocks, as in Figure 6 of Chevigny *et al*. 2022.

**Table 1.**
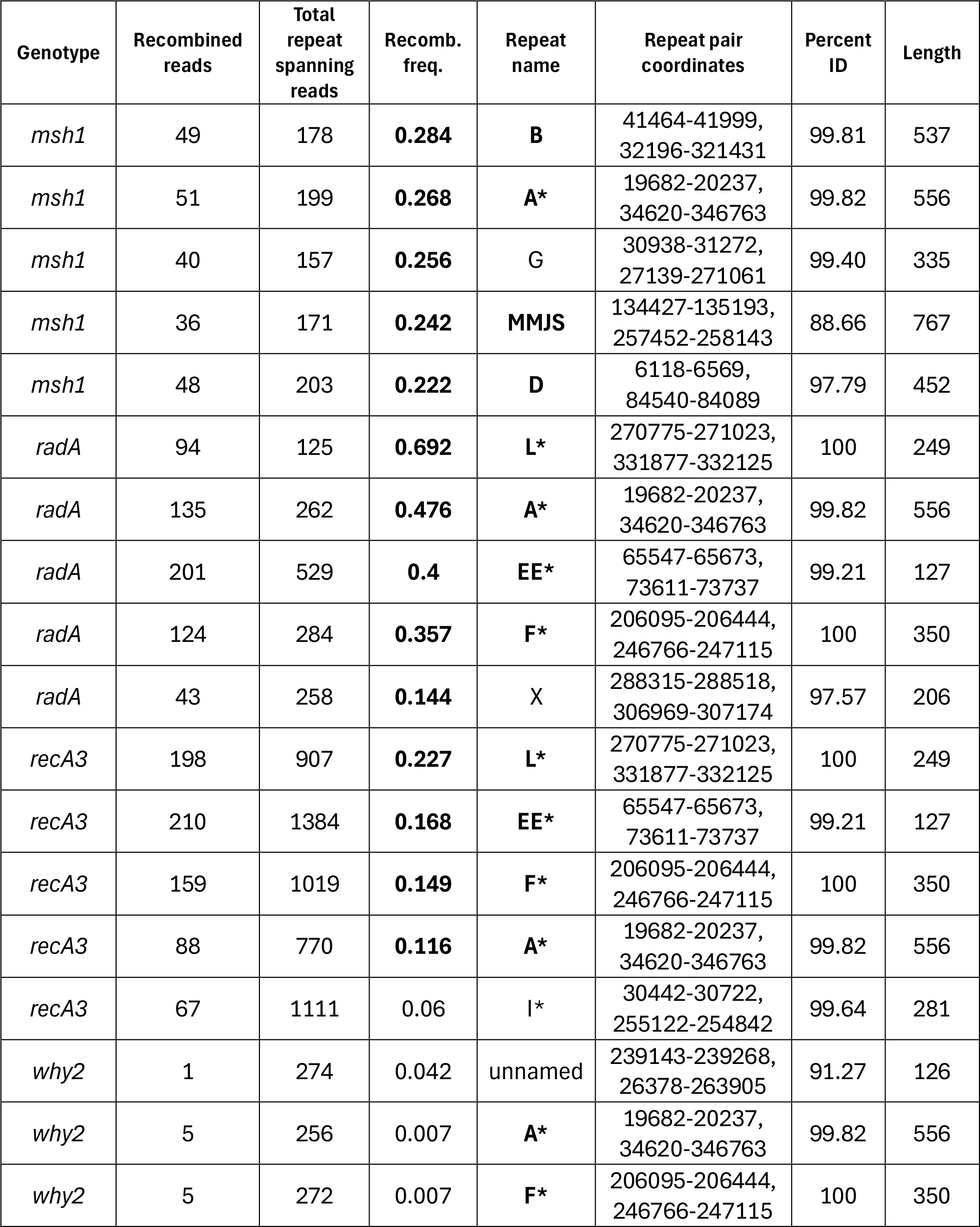
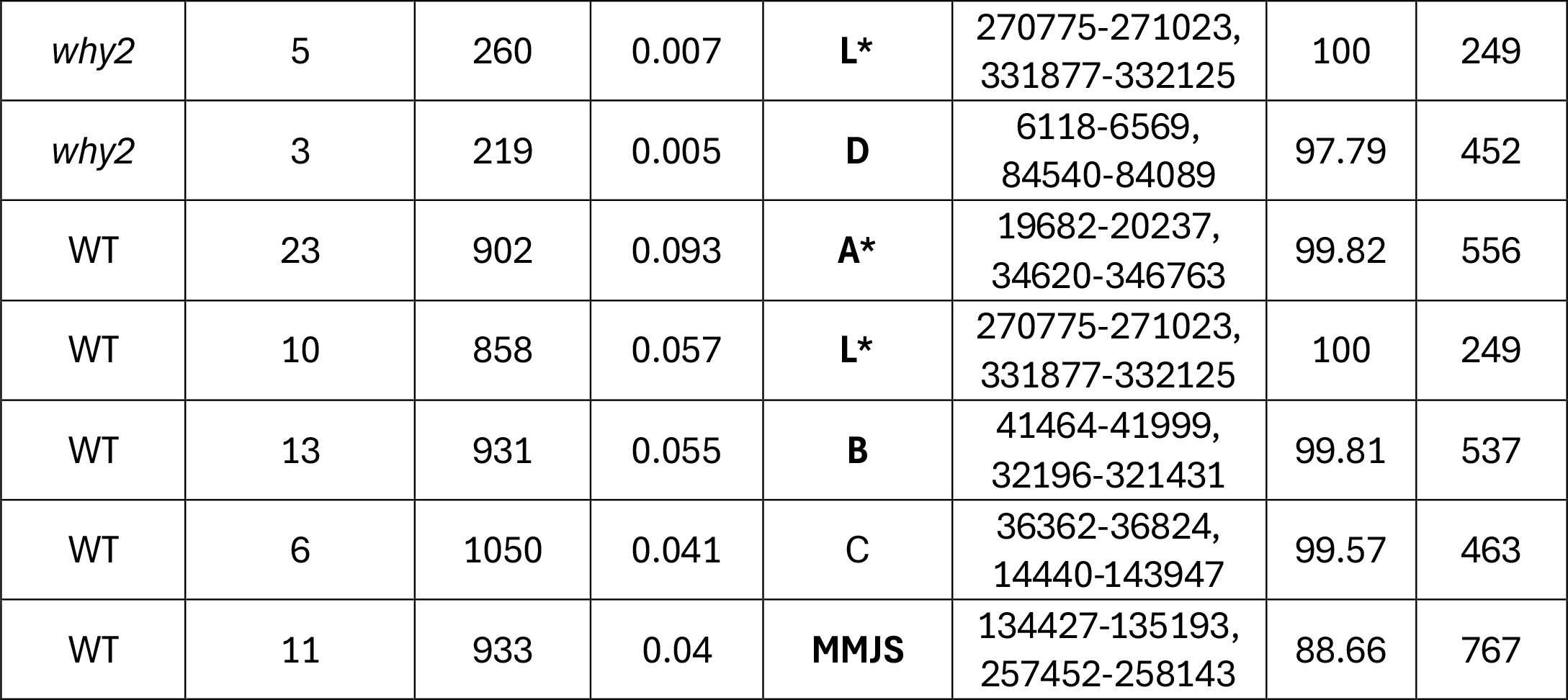
Repeat-specific recombination frequencies at the five most recombinationally active mtDNA repeats for each genotype Listed are the five most active repeats for each genotype, ordered by the recombination frequency within each genotype. Repeat names were sourced from Table S11 of Zou *et al*., 2022. For the *msh1* mtDNA analysis, we relied exclusively the plastid-derived *msh1* samples, and for the *radA* mtDNA analysis, we used a combination of the low coverage *radA* mitochondrial samples and the plastid *radA* samples (see main text). For the WT comparison, we took the average across the single matched WT libraries that were sequenced with each mutant line, including *msh1*and *radA* WT plastid samples (Table S4). The repeats that are also plotted in Figure 8 are denoted with an asterisk. Repeats which make are among the top five most active repeats in more than one genotype are bolded. Repeat-specific recombination frequencies that exceed 0.1 are shown in bold, and note that none of the WT or *why2* repeat specific recombination frequencies meet this threshold.

| Genotype | Recombined reads | Total repeat spanning reads | Recomb. freq. | Repeat name | Repeat pair coordinates | Percent ID | Length |
| --- | --- | --- | --- | --- | --- | --- | --- |
| *msh1* | 49 | 178 | **0.284** | **B** | 41464-41999,<br>32196-321431 | 99.81 | 537 |
| *msh1* | 51 | 199 | **0.268** | **A\*** | 19682-20237,<br>34620-346763 | 99.82 | 556 |
| *msh1* | 40 | 157 | **0.256** | **G** | 30938-31272,<br>27139-271061 | 99.40 | 335 |
| *msh1* | 36 | 171 | **0.242** | **MMJS** | 134427-135193,<br>257452-258143 | 88.66 | 767 |
| *msh1* | 48 | 203 | **0.222** | **D** | 6118-6569,<br>84540-84089 | 97.79 | 452 |
| *radA* | 94 | 125 | **0.692** | **L\*** | 270775-271023,<br>331877-332125 | 100 | 249 |
| *radA* | 135 | 262 | **0.476** | **A\*** | 19682-20237,<br>34620-346763 | 99.82 | 556 |
| *radA* | 201 | 529 | **0.4** | **EE\*** | 65547-65673,<br>73611-73737 | 99.21 | 127 |
| *radA* | 124 | 284 | **0.357** | **F\*** | 206095-206444,<br>246766-247115 | 100 | 350 |
| *radA* | 43 | 258 | **0.144** | **X** | 288315-288518,<br>306969-307174 | 97.57 | 206 |
| *recA3* | 198 | 907 | **0.227** | **L\*** | 270775-271023,<br>331877-332125 | 100 | 249 |
| *recA3* | 210 | 1384 | **0.168** | **EE\*** | 65547-65673,<br>73611-73737 | 99.21 | 127 |
| *recA3* | 159 | 1019 | **0.149** | **F\*** | 206095-206444,<br>246766-247115 | 100 | 350 |
| *recA3* | 88 | 770 | **0.116** | **A\*** | 19682-20237,<br>34620-346763 | 99.82 | 556 |
| *recA3* | 67 | 1111 | 0.06 | **I\*** | 30442-30722,<br>255122-254842 | 99.64 | 281 |
| *why2* | 1 | 274 | 0.042 | unnamed | 239143-239268,<br>26378-263905 | 91.27 | 126 |
| *why2* | 5 | 256 | 0.007 | **A\*** | 19682-20237,<br>34620-346763 | 99.82 | 556 |
| *why2* | 5 | 272 | 0.007 | **F\*** | 206095-206444,<br>246766-247115 | 100 | 350 |
| *why2* | 5 | 260 | 0.007 | **L\*** | 270775-271023,<br>331877-332125 | 100 | 249 |
| *why2* | 3 | 219 | 0.005 | **D** | 6118-6569,<br>84540-84089 | 97.79 | 452 |
| WT | 23 | 902 | 0.093 | **A\*** | 19682-20237,<br>34620-346763 | 99.82 | 556 |
| WT | 10 | 858 | 0.057 | **L\*** | 270775-271023,<br>331877-332125 | 100 | 249 |
| WT | 13 | 931 | 0.055 | **B** | 41464-41999,<br>32196-321431 | 99.81 | 537 |
| WT | 6 | 1050 | 0.041 | C | 36362-36824,<br>14440-143947 | 99.57 | 463 |
| WT | 11 | 933 | 0.04 | **MMJS** | 134427-135193,<br>257452-258143 | 88.66 | 767 |

We analyzed the nanopore reads for evidence of repeat-mediated recombination. To do so, we calculated recombination frequencies for each repeat pair as the count of nanopore reads that recombined at a given repeat (according the BLASTn alignments generated by HiFiSr (Zou *et al*. 2022)) divided by the total number of reads that mapped to the repeat. Table 1 shows the five repeats with the highest recombination frequency for each mutant genotype and the matched WT controls. Fig. 9 shows examples of how the long nanopore reads map to the mitochondrial genome following recombination at inverted (Fig. 9A) or directed repeats (Fig. 9B and C).

**Figure 9.**
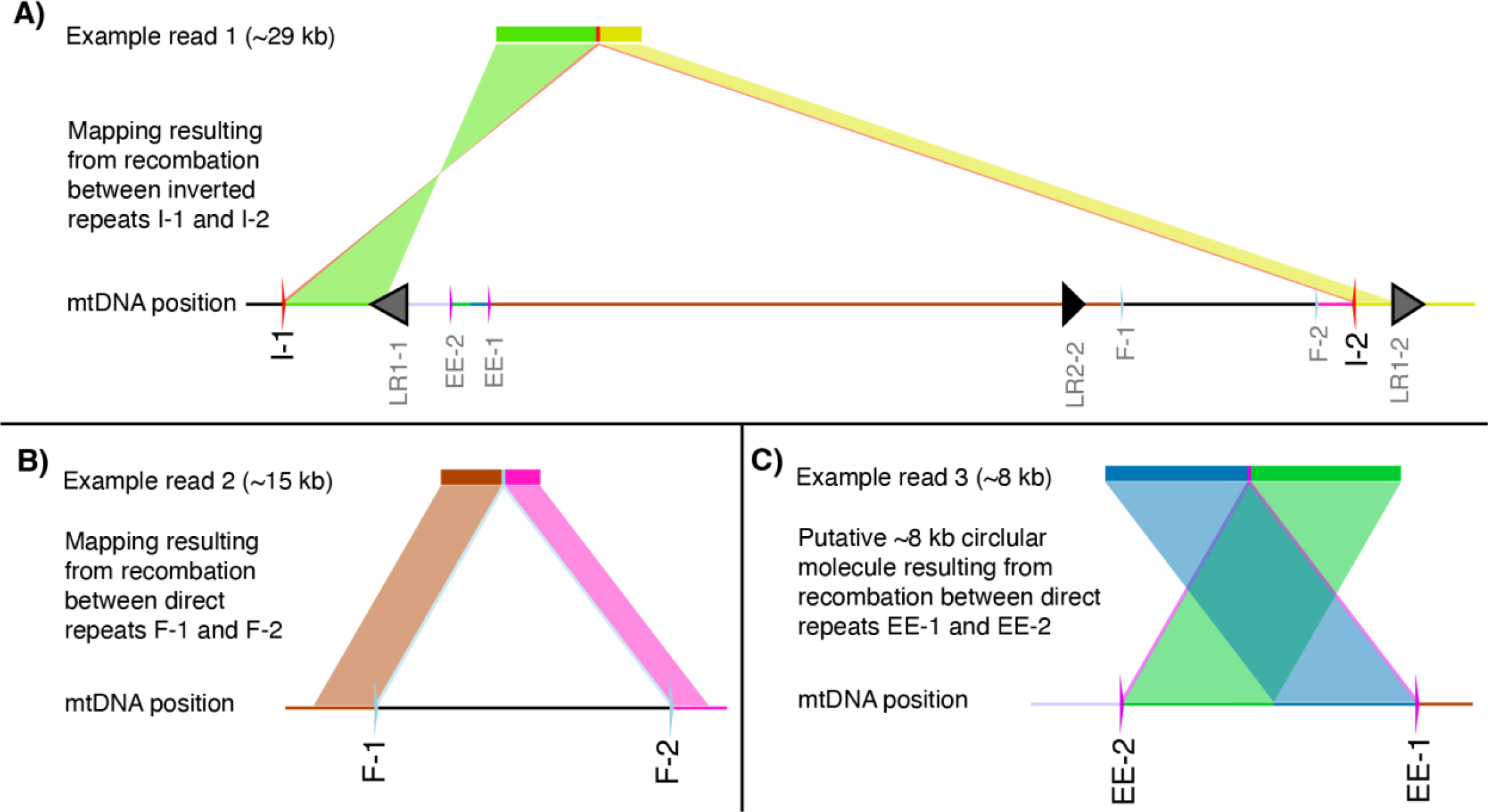
Examples of 3 nanopore reads from *radA* mitochondrial replicate 1 that capture repeat-mediated recombination. Nanopore reads that derive from recombination between inverted repeats map with two hits, one in the forward orientation and the other in the reverse orientation, both flanked by the sequence of a repeat, as shown in A where the 29- kb read is flanked by repeats I-1 and I-2. Recombination between direct repeats results in two hits in the same orientation with a deletion of the intervening sequence (B). The alternative product of recombination between direct repeats is the production of a small circular molecule. We identified a number of putative circular molecules or tandem duplications mediated by recombination between repeats EE-1 and EE-2, which map with two hits in the same orientation, but with a section of the end of the read mapping in front of the end of the read (C).

We calculated genome-wide recombination frequencies for the mtDNA by summing across repeats with at least 10 recombining reads (File S2). The threshold was lowered to repeats with at least three recombining reads in the cpDNA given the smaller number of recombining reads observed in the cpDNA (File S3). We found significant differences in the frequency of mtDNA rearrangements among the WT and mutant lines (one-way ANOVA, p = 1.5×10^-8^, Fig. 10), which were driven by increases in recombination frequency in *msh1, radA* and *recA3* compared to WT (Tukey pairwise comparison, p = 3.0×10^-7^, 2.0×10^-7^, and 0.02, respectively). In contrast, there was no mtDNA recombination frequency difference between *why2* mutants and WT samples (Tukey pairwise comparison, p = 0.99). We found that different repeats apparently become active in different mutant background as evidenced by a two-way ANOVA with a significant interaction between genotype and repeat (p < 2.0×10^-16^). Because our analysis focuses on reads with two or fewer BLASTn hits, we may have underestimated global recombination frequencies, especially in mutant backgrounds, as a PacBio HiFi study found that such reads with three or more BLASTn hits (which arise when reads span two or more repats that have recombined) comprise 0.34% and 8.69% of all reads in WT and *msh1*, respectively (Zou *et al*. 2022). Consistent with previous characterization of repeat mediated recombination in plant mtDNAs (Arrieta-Montiel *et al*. 2009; Davila *et al*. 2011; Miller-Messmer *et al*. 2012; Chevigny *et al*. 2022; Zou *et al*. 2022), we found that repeat length and percent identity are also predictive of recombination frequency through a three-way ANCOVA with repeat length and percent identity as continuous variables (p = 1.8×10^-12^ and 1.4×10^-6^, respectively) and genotype as a categorical variable (p = 2.0×10^-22^). There were no significant differences in repeat-mediated recombination between any of the cpDNA mutants (*msh1, radA, osb2,* and *recA1*) compared to the WT samples (one-way ANOVA, p = 0.849; Fig. 9). We identified no insertions in the HiFiSr variant calls (after requiring at least two nanopore reads to support a putative insertion) and only a single cpDNA deletion of 106 bp in *msh1* mutant replicate 2, which was supported by 18 independent nanopore reads (cpDNA position 148490- 148596).

**Figure 10.**
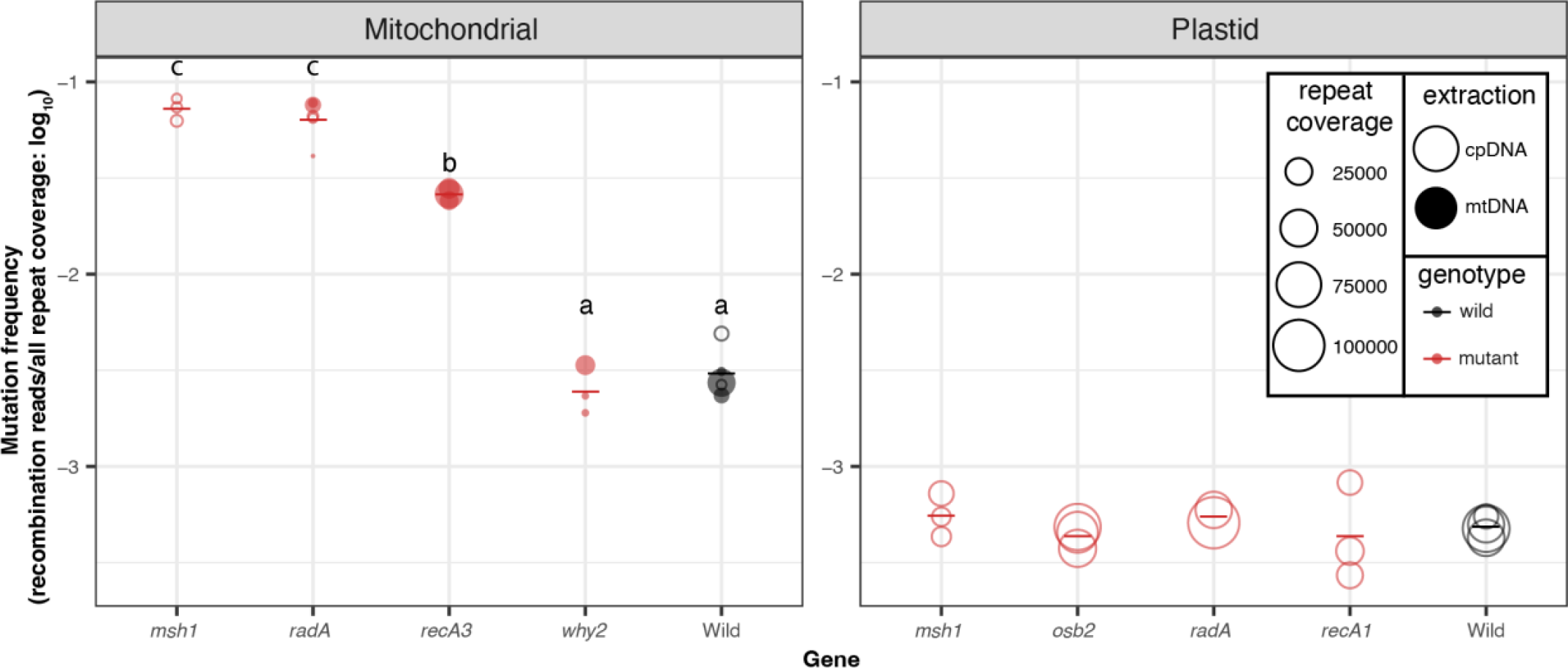
Frequency of repeat-mediated structural variants in the nanopore data. The individual biological replicates are plotted as circles with the size of the circle scaled by the number of repeats that are covered in the nanopore alignments. Closed circles are the libraries from mitochondrial extractions, while the open circles are libraries from the plastid extractions. In some cases, cpDNA extractions were used to harvest contaminating mtDNA-mapping reads because of low yield from direct sequencing of the mtDNA extractions. Group averages are plotted as dashes. Mutants are plotted in red, while WT samples are plotted in black. Letters represent statistically significant groupings according to Tukey pairwise comparisons on a one-way ANOVA (p<0.001). There were no differences among plastid genotypes.

## DISCUSSION

### Potential causes of elevated organellar mutation rates in lines with disrupted recombination machinery

By utilizing highly accurate Duplex Sequencing for point mutation detection and long-read Oxford Nanopore sequencing for structural variant detection, we have characterized the overall organellar mutational dynamics in *A. thaliana* lines lacking genes with roles in organellar genome recombination. The increases in point mutations we observed in *radA, recA3,* and *recA1* are much smaller than the effects previously observed in *msh1* mutants (Wu *et al*. 2020) where mutants experience 6.0-fold and 116.5-fold increases in SNVs (in mtDNA and cpDNA, respectively) and 86.6-fold and 790.6-fold increases in indels (in mtDNA and cpDNA, respectively). In contrast, *radA* mutants incurred 2.6-fold and 12.6-fold more mtDNA and cpDNA SNVs (respectively) and 5.1-fold and 3.1-fold more mtDNA and cpDNA indels (respectively) than the matched WT controls. The point mutation increases in *recA3* and *recA1* were even smaller than in the *radA* mutants. One complication with directly comparing the mutant vs. WT fold changes across the newly generated mutant lines compared to those generated in Wu *et. al.,* (2020) is the decrease in WT mutation rates in the new genes (Fig. 2). Because of the shift in the baseline WT rates, the numbers cited above may actually underestimate the gap in effect size between *msh1* and the newly analyzed genes.

The point mutation increases in *msh1* mutants have clear mechanistic explanations which were first predicted based on the MSH1 mismatch recognition and GIY-YIG endonuclease domains (Christensen 2014; Wu *et al*. 2020). In contrast, given that RADA, RECA3 and RECA1 are all thought to function in the resolution of recombination intermediates, it is more difficult to explain the mechanisms responsible for increased point mutations in these lines. One possibility is that in the absence of one recombination pathway, recombining molecules are shuttled into an alternative, less faithful recombination pathway. For example, in mutant lines deficient in homologous recombination (HR), double-stranded breaks (DSBs) may be repaired via error prone non-homologous end joining (NHEJ) or MMEJ, which could drive increases in indels and SNVs (Waters *et al*. 2014; García-Medel *et al*. 2019). Evidence suggests that RADA functions as the principal branch migration factor in a primary mtDNA and cpDNA homologous recombination (HR) pathway, while RECA3 may fill the same role as RADA in a partially redundant and less utilized mtDNA specific-HR pathway (Chevigny *et al*. 2022). Interestingly, RECA2 is thought to initiate recombination in both pathways and is essential in plants (Miller-Messmer *et al*. 2012; Chevigny *et al*. 2022). The larger SNV and indel increases in the *radA* mutants than in the *recA3* mutants may reflect the relative utilization (and importance) of these two partially redundant HR pathways (Chevigny *et al*. 2022).

Similarly, previous studies have documented increased NHEJ and MMEJ in cpDNA of *recA1* mutants (Zampini *et al*. 2015), which is consistent with the significant increase in indels and marginally significant increase in SNVs reported here (Fig. 1).

Another possibility is that the rise in point mutations is an indirect effect of increased repeat-mediated recombination and its associated harm to organelle function. Increased recombination between short repeat sequences may disrupt genes, organellar genome stoichiometry, and genome organellar replication, which is recombination-dependent in plants (Shedge *et al*. 2007; Rowan *et al*. 2010; Chevigny *et al*. 2020). Plant organellar genomes encode proteins necessary for the electron transport chains of respiration and photosynthesis and disruption of these pathways can result in the excess production of DNA damaging reactive oxygen species (ROS; Liu *et al*. 2021). Although a direct link between ROS-mediated damage to DNA and mutation rates remains contentious (Kennedy *et al*. 2013; Itsara *et al*. 2014; Broz *et al*. 2021; Waneka *et al*. 2021; Sanchez-Contreras *et al*. 2021), ROS molecules have been shown to indirectly affect point mutation rates by impairing proofreading capabilities via damage to the metazoan mtDNA polymerase (Pol γ; Anderson *et al*. 2020). Impairment of organellar function is also consistent with phenotypic growth defects in *radA*, which include retarded development and distorted leaves with chlorotic sectors (Chevigny *et al*. 2022).

### Potential explanations of mutational biases based on DNA strand asymmetry and flanking nucleotides

We found that SNVs in the *msh1* mutants and WT plants from Wu *et al.,* (2020) had biased distributions in terms of strand (non-template vs. template) and trinucleotide context.

Such patterns are useful for understanding the underlying mechanisms driving mutation formation (Haradhvala *et al*. 2016; Sun *et al*. 2018; Moeckel *et al*. 2023). For example, CG→TA strand asymmetries documented in diverse metazoan mtDNAs have been proposed to result from the two DNA strands experiencing unequal time in single-stranded states during mtDNA replication, since single-stranded DNA is more vulnerable to cytosine deamination (a primary driver of CG→TA transitions) (Kennedy *et al*. 2013; Itsara *et al*. 2014; Arbeithuber *et al*. 2020; Waneka *et al*. 2021; Sanchez-Contreras *et al*. 2021). In mammals, C→T substitutions are ∼10-fold more common than G→A substitution on the mtDNA heavy strand (H-strand), which likely spends more time in a single-stranded state as the mtDNA is copied via a strand-asynchronous replication mechanism (Kennedy *et al*. 2013; Arbeithuber *et al*. 2020). Further, the C→T substitutions form two gradients starting at the two H-strand origins of replication, consistent with the regions closest to the origin being single stranded for longer (Sanchez-Contreras *et al*. 2021).

The substantial CG→TA strand asymmetries we observed in the mtDNA of the Wu *et al.,* (2020) WT libraries are unlikely to be explained by replication mechanisms given that plants mtDNAs lack discrete origins of replication or dedicated ‘leading and lagging’ strands (alternatively referred to as light and heavy strands, respectively, in some systems) and instead rely on recombination-mediated replication (Gualberto and Newton 2017; Brieba 2019; Chevigny *et al*. 2020). Instead, our strand asymmetry analysis focused on genic regions, motivated by well-established patterns of more C→T than G→A substitution on non-template strands which spend more time in exposed single-stranded during transcription (Haradhvala *et al*. 2016; Vöhringer *et al*. 2021; Moeckel *et al*. 2023).

Surprisingly, we found an opposite pattern with template strands exhibiting far more C→T than G→A substitutions (Fig. 4). This effect was especially pronounced in rRNA and tRNA genes where the C→T substitutions occurred on the template strand in all 32 observed CG→TA transitions. An enrichment of C→T substitutions on template strands also occurred in the mtDNA (but not the cpDNA) of the *msh1* mutants, though there was less power for detecting statistically significant effects (Fig. 5). The overabundance of A→G compared to T→C substitutions in *msh1* mutant cpDNA template strands also occurs in the opposite direction of predicted effects given that the non-template strand is again expected to experience increased adenine deamination (which leads to A→G substitutions; Mugal *et al*. 2009; Sanchez-Contreras *et al*. 2021).

Enrichment of C→T and A→G substitutions on template strands is puzzling, and to our knowledge there are no other instances where this widespread transcriptional asymmetry has been reversed (Mugal *et al*. 2009; Moeckel *et al*. 2023). Reversals in strand asymmetries have been reported in metazoan mitochondrial genomes, but in these cases the asymmetries are replication based, and the reversals are proceeded by an inversion of the origin of replication, effectively switching the leading and lagging strands (Wei *et al*. 2010). It is notable that the WT CG→TA asymmetries are most pronounced in the rRNA and tRNA genes (Fig. 4), which are likely more highly expressed than the protein coding genes. Increases in transcription have been shown to drive genomic instability in the *A. thaliana* cpDNA due to the increased formation of R-loops (RNA/DNA hybrids formed by displacement of the other DNA strand), which stall replication forks and lead to DSBs (Pérez Di Giorgio *et al*. 2019). It is possible that increased mtDNA expression also leads to the formation of R-loops and DSBs which may then be repaired through error prone NHEJ and MMEJ. However, it is not clear how this would drive strand asymmetric mutation.

Further, such a mechanism is not consistent with the relatively even distribution of SNVs across intergenic vs. transcribed regions of the genome (Fig. 3). The magnitude of the CG→TA asymmetries is decreased in the *msh1* mutants (roughly 2-fold averaging across all genic sequences) compared to in the WT controls (roughly 6-fold). This shift may reflect a larger proportional contribution of mutations from simple DNA polymerase misincorporation errors (which are not expected to be strand-biased) in the absence of MSH1 activity.

The CG→TA transitions in the WT lines and both transitions in the *msh1* mutants were also impacted by the identity of neighboring nucleotides (Figs. 6 and 7). Trinucleotide effects have previously been implicated to bias mutation distribution in the *A. thaliana* nuclear genome (Lu *et al*. 2021) as well as in the mtDNAs of various metazoans (Itsara *et al*. 2014; Arbeithuber *et al*. 2020; Waneka *et al*. 2021; Sanchez-Contreras *et al*. 2021). It is noteworthy that the specific trinucleotides associated with CG→TA transitions differ between WT and *msh1* mutants. The 5′ YCN signature (where Y is any pyrimidine and N is any nucleotide) in the WT lines is similar to that induced by APOBEC3-mediated cytosine deamination in human cell lines (Carpenter *et al*. 2023), though plants lack APOBEC enzymes so the relevance of this shared pattern is unclear. Meanwhile, the 5′ NCG signature in the *msh1* mutants is consistent with spontaneous water mediated cytosine deamination (Carpenter *et al*. 2023).

### Patterns of repeat-mediated recombination differs among mutant lines

The repeat mediated mtDNA recombination activity we documented in the *msh1*, *radA* and *recA3* mutants is consistent with the previously documented recombination increases of these mutant backgrounds (Shedge *et al*. 2007; Arrieta-Montiel *et al*. 2009; Rowan *et al*. 2010; Davila *et al*. 2011; Miller-Messmer *et al*. 2012; Zampini *et al*. 2015; Wu *et al*. 2020; Chevigny *et al*. 2022; Zou *et al*. 2022). The absence of an effect in the *why2* mutants is interesting given that *why2* is the most abundant protein in mitochondrial nucleoids (Fuchs *et al*. 2020) and plants lacking *why2* display aberrant mitochondrial morphology (Golin *et al*. 2020; Negroni *et al*. 2024). On the other hand, this result is consistent with a previous study that showed *why2* mutants become more recombinationally active than WT under increased genotoxic stress (ciprofloxacin treatment) but showed no recombinational difference from WT under ‘normal’ growth conditions (Cappadocia *et al*. 2010; Negroni *et al*. 2024).

Though *msh1*, *radA* and *reca3* are all required for the suppression of repeat-mediated recombination in mtDNA, these proteins likely function either in independent HR pathways (*radA, recA3*) or in different ways (*msh1*). As, noted, RECA3 is thought to facilitate branch migration in an HR pathway that may be relatively minor compared to the one in which RADA functions (Chevigny *et al*. 2022). Previous studies of *recA3/msh1* and *recA3/radA* double mutants have shown the double mutants are more recombinationally active than *recA3* single mutants (Shedge *et al*. 2007), supporting the hypothesis that *RECA3-*mediated HR is at least partially independent of *RADA*-mediated HR (Miller-Messmer *et al*. 2012; Chevigny *et al*. 2022). This model is supported by the greater increase in global recombination frequency in *radA* compared to *recA3* (Fig. 10). We might also expect different repeats to become active in *recA3* compared to *radA* mutants. However, as seen in Table1, there is substantial overlap in the repeats with increased recombination frequencies in these mutants, though the extremely high recombination frequency at repeat L in *radA* is one major difference. Meanwhile, MSH1 has been proposed to suppress non-allelic recombination by recognizing and rejecting mismatches in the invading strand during heteroduplex formation (Christensen 2018; Broz *et al*. 2022), which could be a shared feature in both *RADA* and *RECA3* dependent HR pathways. Supporting this idea, there is an increased number of repeats that become active in *msh1* mutants compared to *radA* and *recA3* mutants. Specifically, there are 12 repeat pairs with a recombination frequency greater than 0.1 in *msh1* mutants but only four and nine repeat pairs that meet this threshold in *recA3* and *radA* mutants, respectively (File S2).

Given that recombination is activated differently between the mutants (Fig. 8), the high degree of repeatability between replicates is fascinating (Figs. S5, S6, S7, S8). These repeatable patterns rely on consistent activation of distinct repeat pairs and/or consistent maintenance/replication of certain recombination products. Understanding why different repeats become active and how these patterns relate to the increase in point mutations reported here remains an important unanswered question in the field of plant organellar genome maintenance.

## DATA AVAILABILITY

The Duplex Sequencing and Oxford Nanopore reads were deposited to the NCBI Sequence Read Archive (SRA) under BioProject PRJNA1113549.

## SUPPLEMENTAL FIGURES

**Figure S1.**
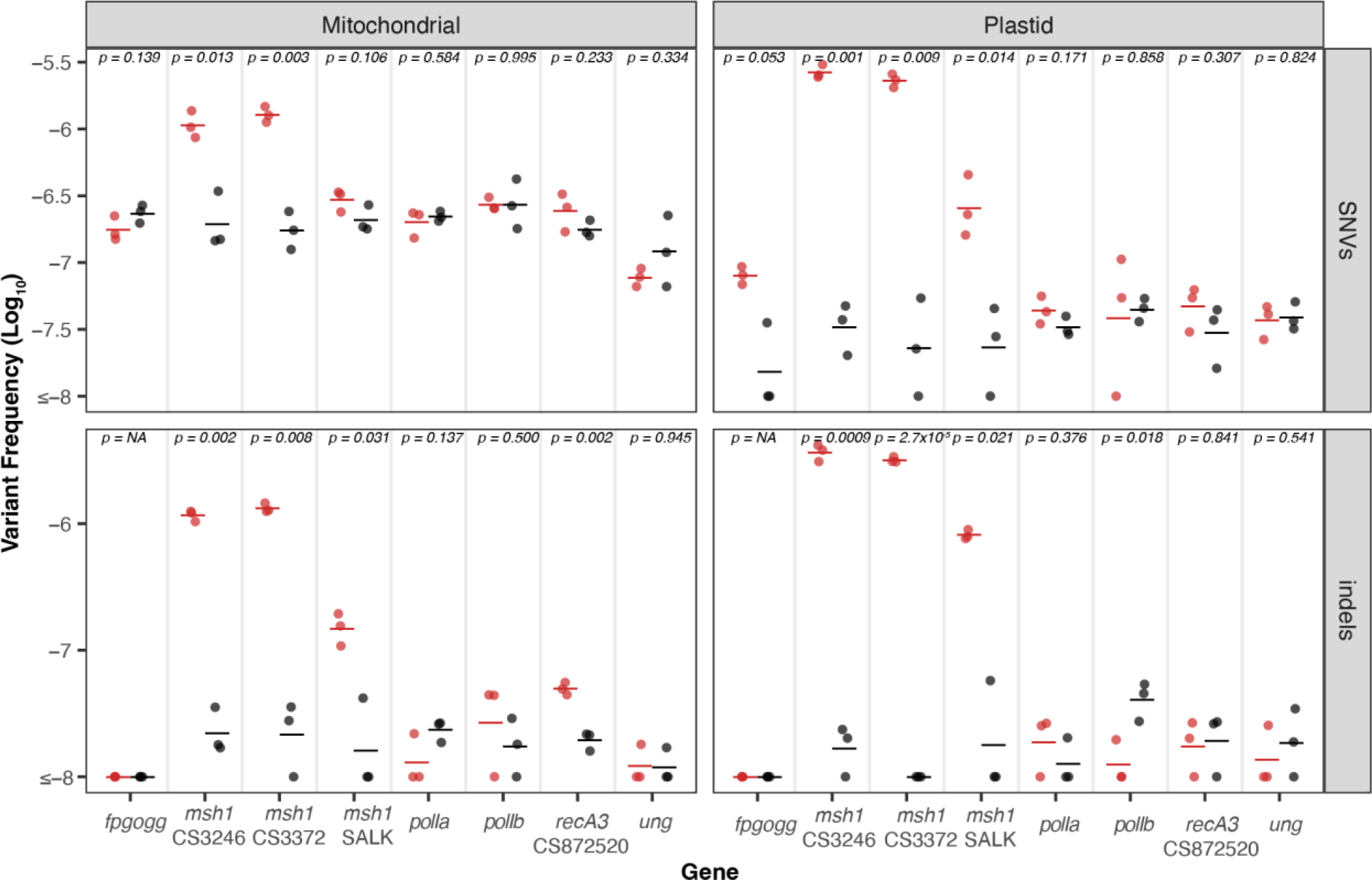
*De novo* point mutations measured with Duplex Sequencing from data generated in Wu *et al*. 2020. For each gene of interest (x-axis) mutant lines are plotted in red and matched WT controls are plotted in black. The individual biological replicates are plotted as circles, and group averages are plotted as dashes. Panels separate the data by genome (left column: Mitochondria and right column: Plastid) and by point mutation type (top row: SNVs and bottom row: indels). The y-axis shows the log-transformed SNV frequencies (total SNVs/total DCS coverage). P-values show the result of a two-tailed *t*-test comparing WT vs mutant mutation frequencies for each gene of interest. We found significant increases in SNV and indel frequencies in the *msh1* CS3246 and *msh1* CS3372 mutants (both genomes) but the *msh1* SALK046763 mutant, which is not a complete knockout of the *msh1* gene (Wu *et al.,* 2020) had weaker effects. In addition, we note that this *recA3* null allele is different from the *recA3* null allele that was reported in the new dataset, but both yielded similar results: significant indel and weakly significant SNV increases in mtDNA of the *recA3* mutant. Also note the marginally significant difference in *fpg*/*ogg* plastid SNVs is explained by just 5 SNVs in mutants and a single SNV in the WT controls, which we do not consider to be a biologically meaningful difference.

**Figure S2.**
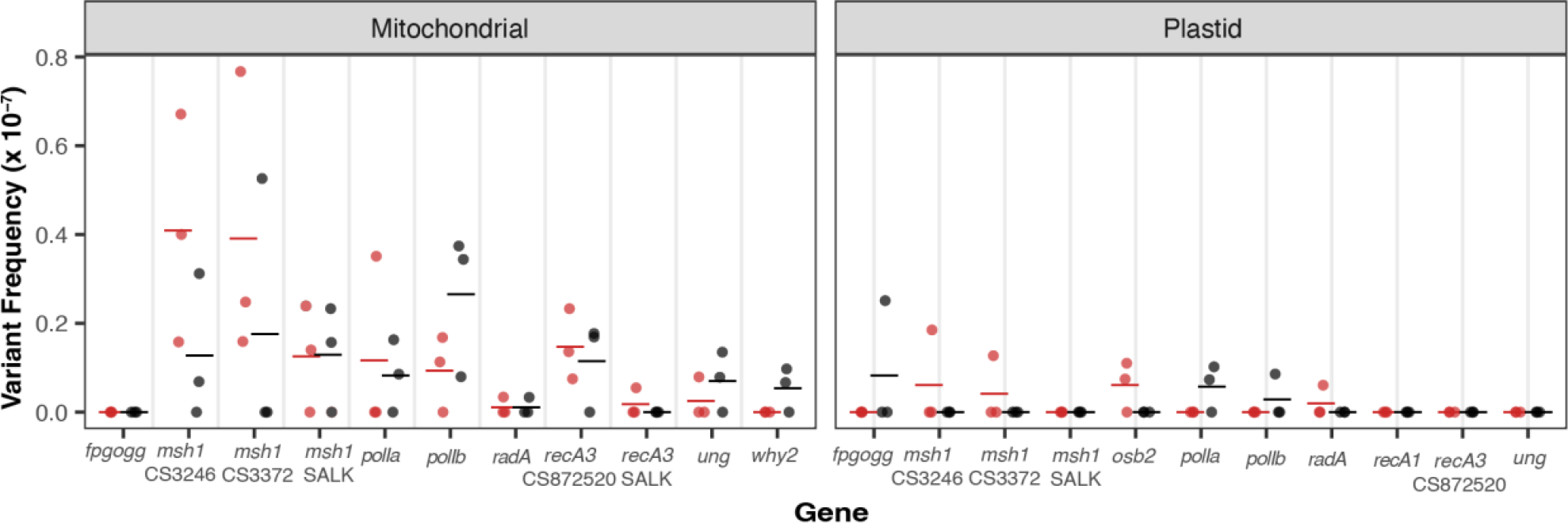
Dinucleotide mutations measured with Duplex Sequencing. For each gene of interest (x axis) mutant lines are plotted in red, and matched WT controls are plotted in black. The individual biological replicates are plotted as circles, and group averages are plotted as dashes. Panels divide the data by mitochondrial and plastid. We performed Wilcoxon rank sum tests to look for differences between mutant and matched WT controls and all p-values were > 0.05. Note that *recA3* CS872520 dataset was generated in Wu *et al*. (2020), and the *recA3* SALK 146388 dataset was generated in this study.

**Figure S3.**
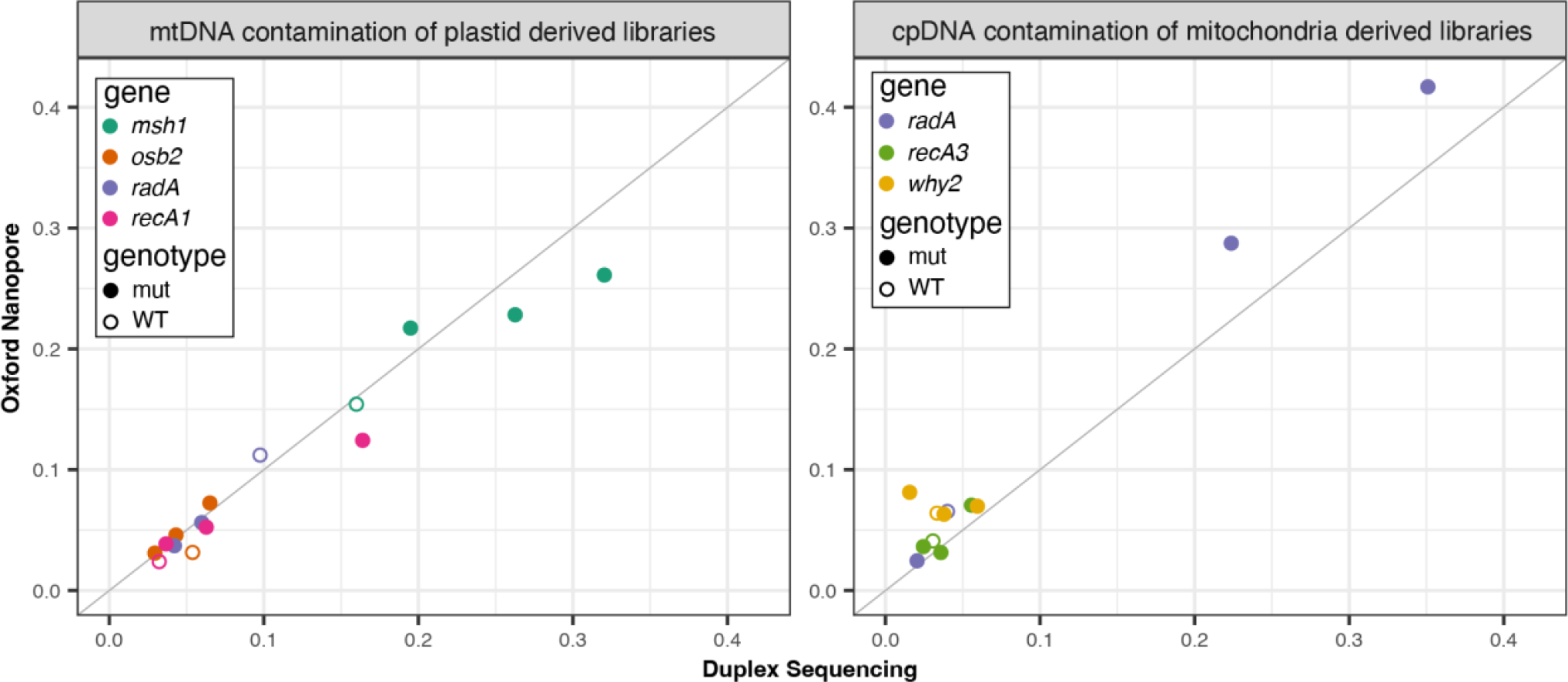
Correlation of cross-organelle contamination in Oxford Nanopore and Duplex Sequencing libraries. Contamination is calculated as the number of contaminating reads in the read alignments divided by the total number of organellar alignments. The different mutant lines are colored according to the figure legend with mutant replicates plotted using closed circles and matched WT controls plotted with open circles. The 1:1 diagonal line is shown in gray. Though the level of contamination varies between different DNA samples (for example mtDNA contamination is higher in the plastid derived *msh1* libraries) the contamination levels are generally similar irrespective of sequencing technique. Note, For the *msh1* mtDNA analysis, we relied exclusively the plastid-derived *msh1* samples, and for the *radA* mtDNA analysis, we used a combination of the low coverage *radA* mitochondrial samples and the plastid *radA* samples (see main text).

**Figure S4.**
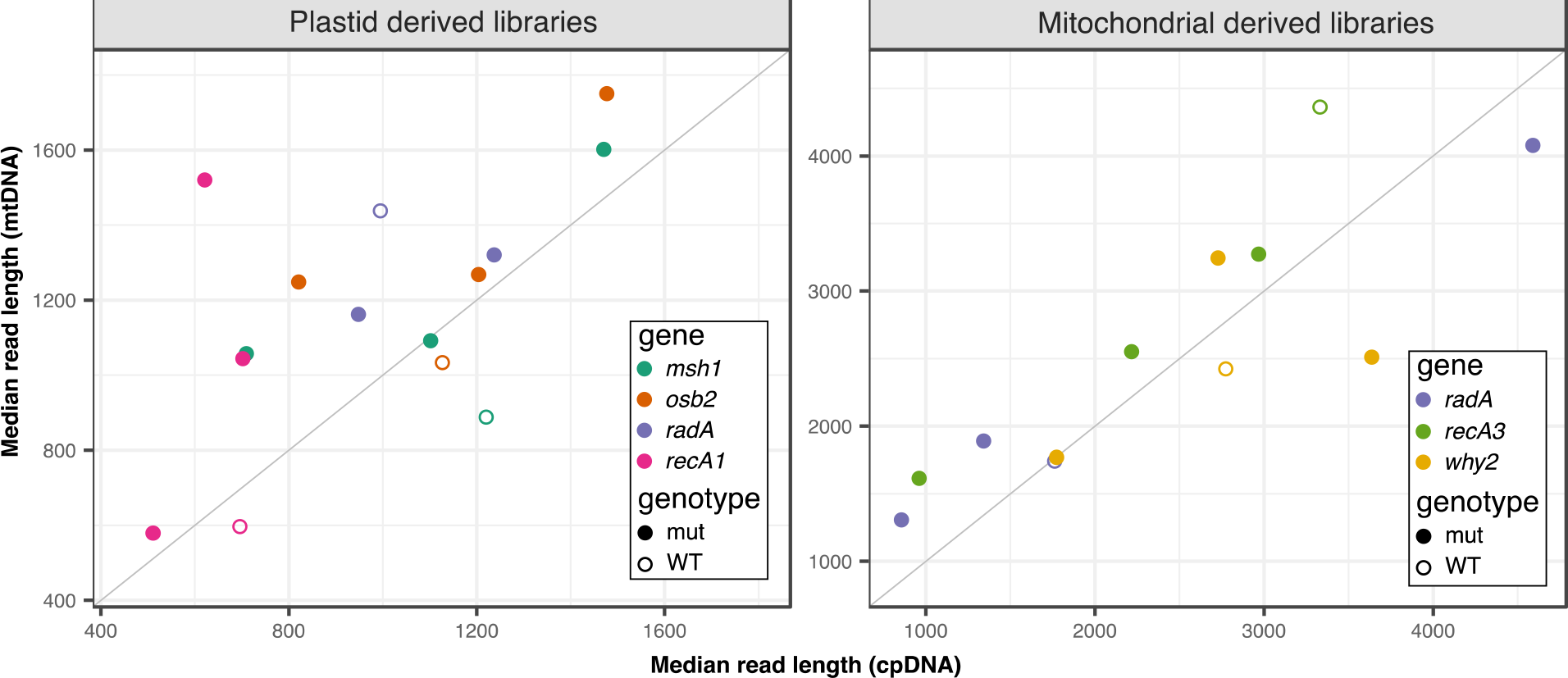
Median read length cross-organelle contaminating and native reads in the plastid and mitochondrial derived nanopore libraries. The different mutant lines are colored according to the figure legend with mutant replicates plotted using closed circles and matched WT controls plotted with open circles. The 1:1 diagonal line is show in gray. Note, For the *msh1* mtDNA analysis, we relied exclusively the plastid-derived *msh1* samples, and for the *radA* mtDNA analysis, we used a combination of the low coverage *radA* mitochondrial samples and the plastid *radA* samples (see main text).

**Figure S5.**
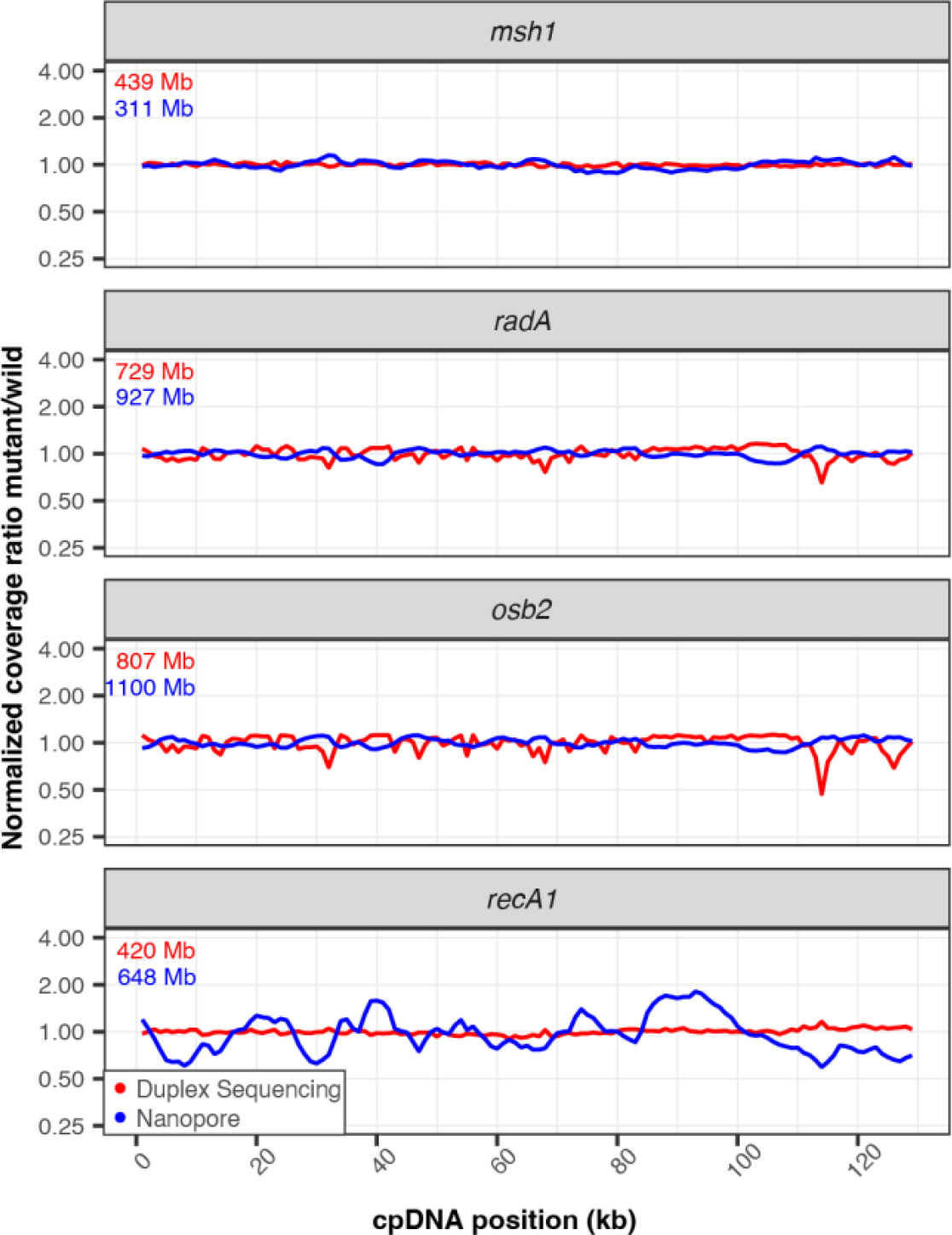
Normalized coverage of plastid genomes in mutant lines of interest . Coverage of each Duplex Sequencing (red) or nanopore (blue) library was calculated in 1000-bp windows. Mutant coverage was pooled and divided by WT coverage and the resulting ratios were normalized to 1 for plotting. The total amount of sequencing data used to generate each plot is shown in the top left corner of each panel (red=Duplex Sequencing and blue=nanopore) and is included to highlight the instances where disagreement between the Duplex Sequencing and nanopore lines may be explained by increased variance in the nanopore sample due to lower mtDNA coverage. To see the coverage of the individual replicates see Fig S8 and S9.

**Figure S6.**
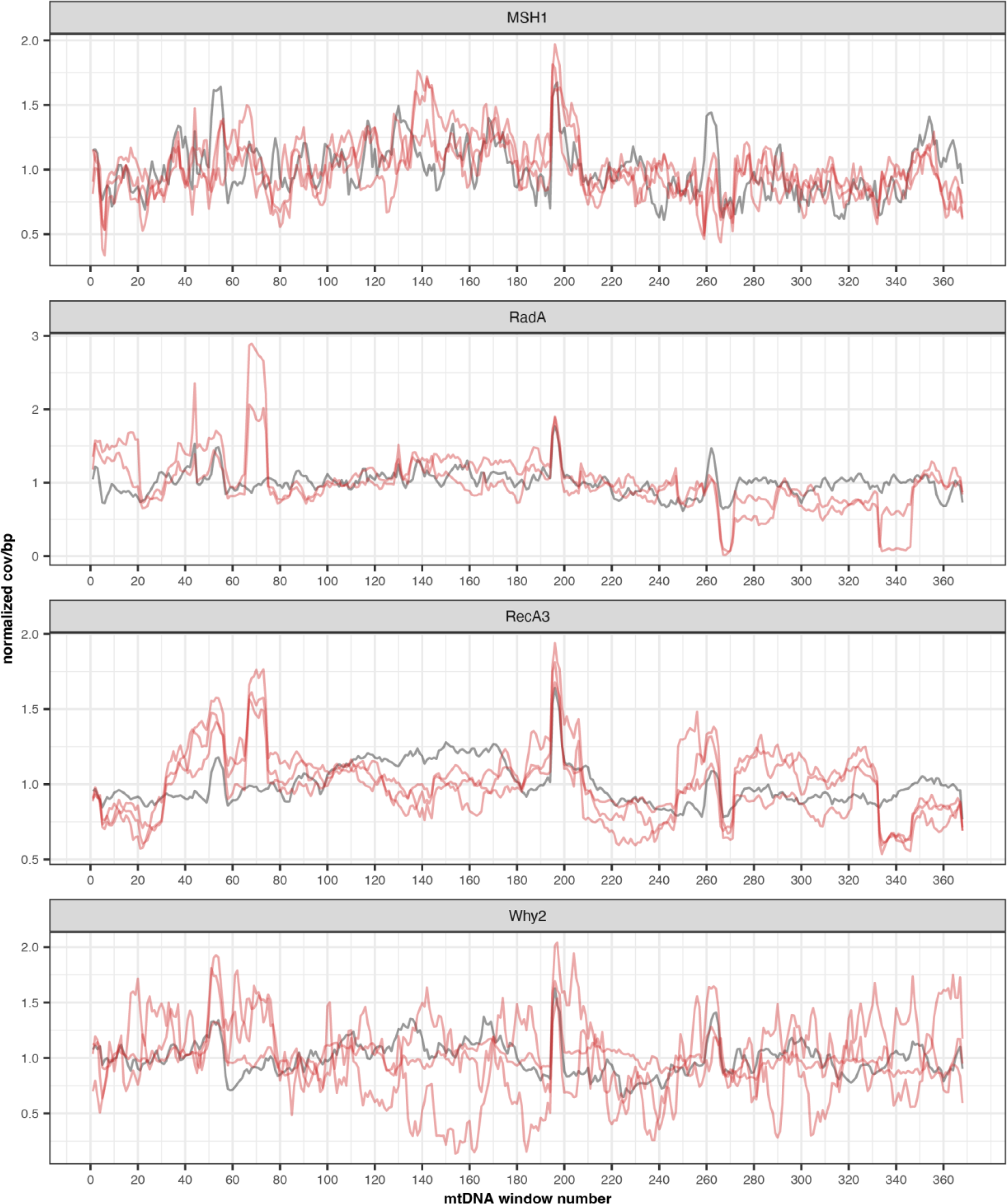
Normalized coverage of the individual nanopore mtDNA replicates (used to generate Fig. 8). The red and black lines show the normalized coverage of the mutant replicates and the matched WT control, respectively. Note that variation in the *why2* mutants is likely due to extremely low coverage in these samples (average coverage per bp of 157.3, 6.5 and 7.0 in mutant replicates 1, 2 and 3, respectively).

**Figure S7.**
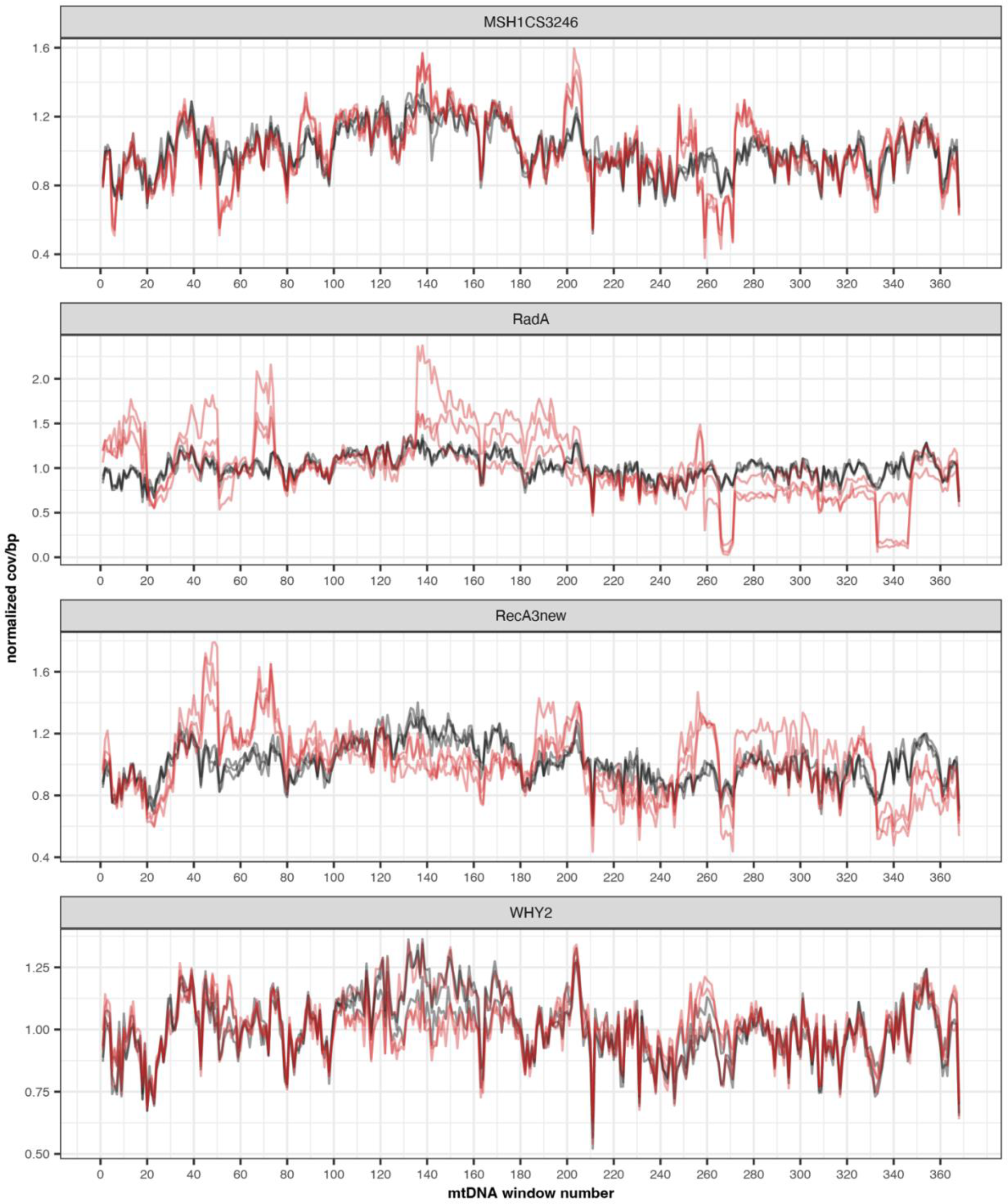
Depth of coverage of the individual Duplex Sequencing mtDNA replicates (used to generate Fig. 8). The red and black lines show the normalized coverage of the mutant replicates and the matched WT control, respectively.

**Figure S8.**
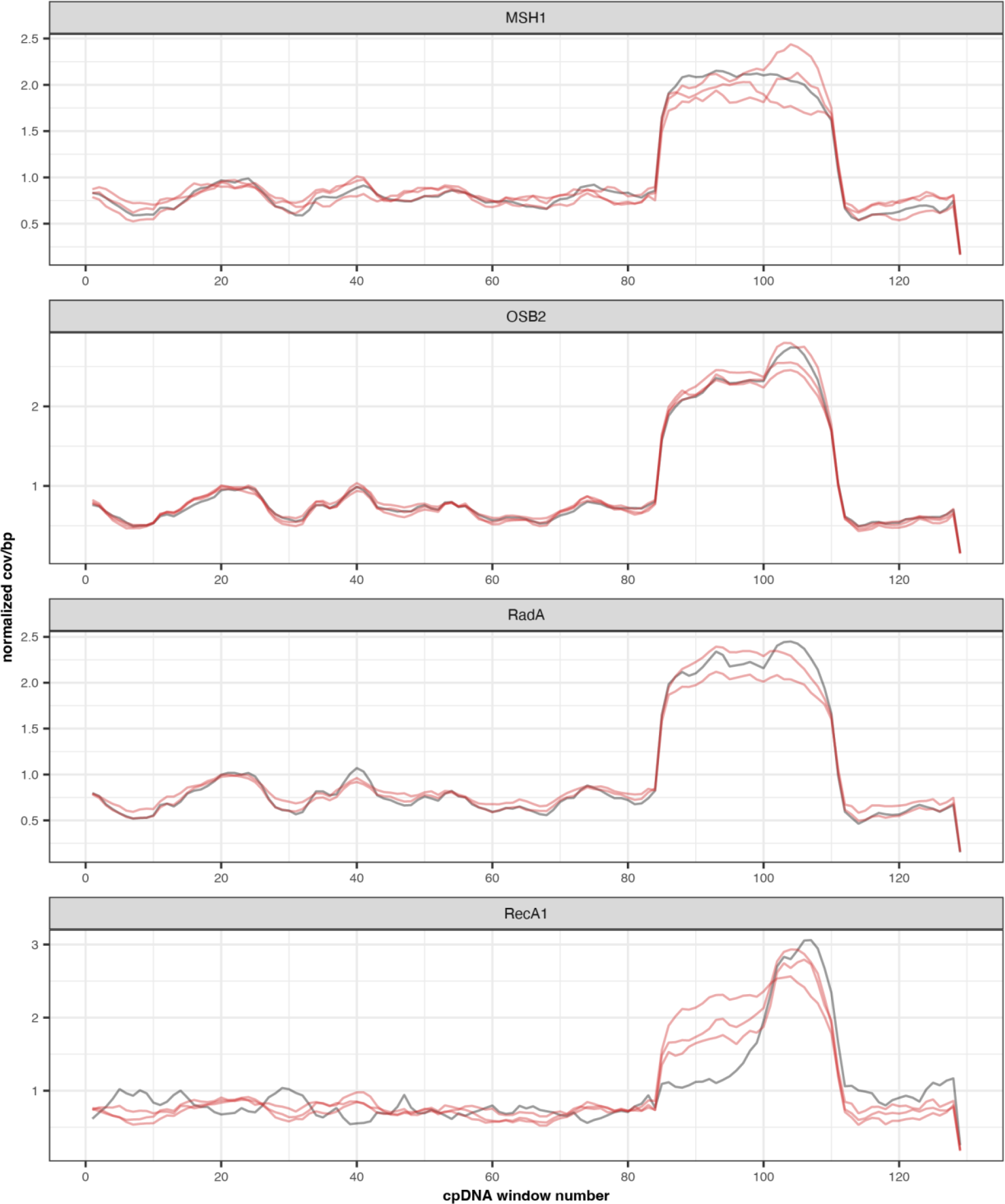
Normalized of coverage of the individual nanopore cpDNA replicates (used to generate Fig. 8). The red and black lines show the normalized coverage of the mutant replicates and the matched WT control, respectively. Note that the spike in coverage at ∼84-112 kb results from the large inverted repeat, since these reads were mapped noncompetitively with minimap2 (see methods). The second copy of the inverted repeat was omitted for plotting.

**Figure S9.**
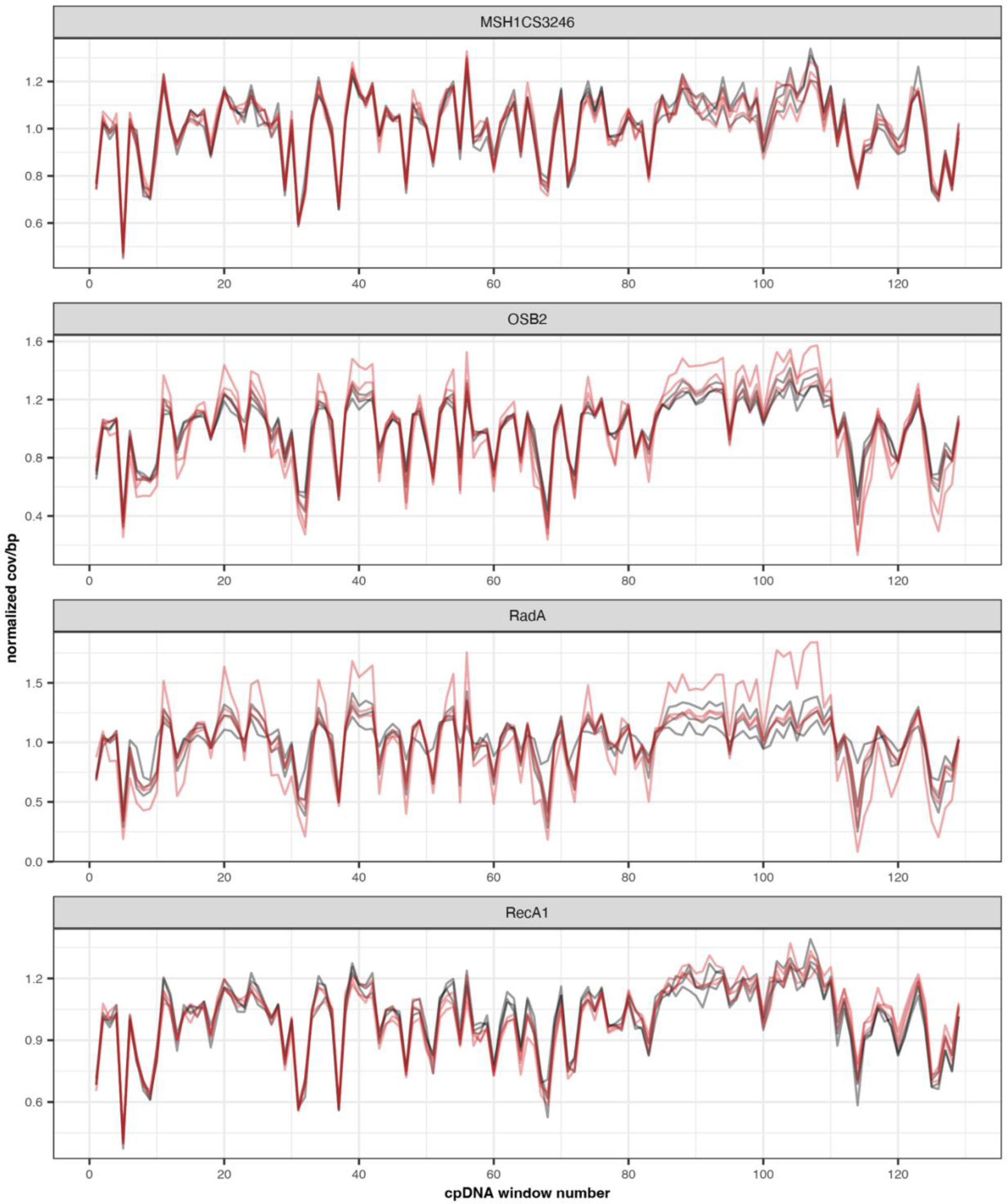
Normalized coverage of the individual Duplex Sequencing cpDNA replicates (used to generate Fig. 8). The red and black lines show the normalized coverage of the mutant replicates and the matched WT control, respectively. Note these reads were mapped to a full length cpDNA but the second large inverted repeat was omitted for plotting.

## SUPPLEMENTAL TABLES

**Table S1.**
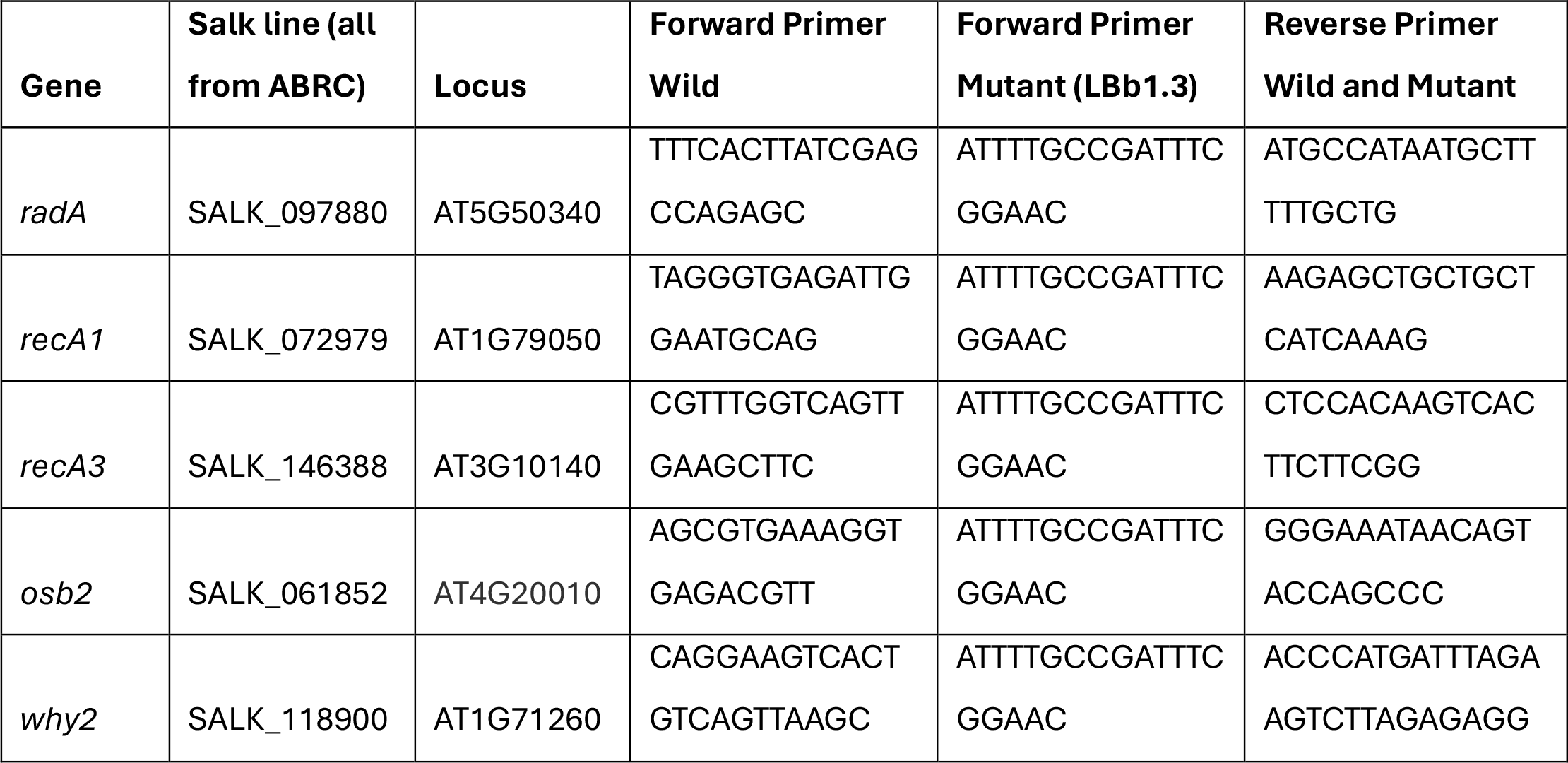
Mutant lines used in this study and primers to verify plant genotype

**Table S2.** Duplex read-pairs and organellar genome coverage

| Sample | Count of read-pairs (2x150) | Organellar coverage per bp |
| --- | --- | --- |
| mitochondrial_rada_mut_1 | 75863350 | 297.9 |
| mitochondrial_rada_mut_2 | 64771627 | 281.8 |
| mitochondrial_rada_mut_3 | 139195192 | 803.4 |
| mitochondrial_rada_wild_1 | 70847671 | 127.8 |
| mitochondrial_rada_wild_2 | 61998713 | 246.9 |
| mitochondrial_rada_wild_3 | 127161861 | 816.4 |
| mitochondrial_reca3_mut_1 | 43472074 | 94.2 |
| mitochondrial_reca3_mut_2 | 44312097 | 229.3 |
| mitochondrial_reca3_mut_3 | 59128403 | 497.3 |
| mitochondrial_reca3_wild_1 | 62354817 | 238.2 |
| mitochondrial_reca3_wild_2 | 54311915 | 183.2 |
| mitochondrial_reca3_wild_3 | 40734051 | 144.8 |
| mitochondrial_why2_mut_1 | 63375069 | 338.0 |
| mitochondrial_why2_mut_2 | 76906783 | 284.9 |
| mitochondrial_why2_mut_3 | 76221972 | 292.6 |
| mitochondrial_why2_wild_1 | 68231709 | 279.8 |
| mitochondrial_why2_wild_2 | 81396138 | 379.9 |
| mitochondrial_why2_wild_3 | 86880259 | 408.4 |
| plastid_osb2_mut_1 | 47505179 | 1176.6 |
| plastid_osb2_mut_2 | 54307516 | 870.8 |
| plastid_osb2_mut_3 | 59415250 | 898.6 |
| plastid_osb2_wild_1 | 59542949 | 1132.8 |
| plastid_osb2_wild_2 | 69408084 | 889.7 |
| plastid_osb2_wild_3 | 67727784 | 668.6 |
| plastid_rada_mut_1 | 76116128 | 1174.4 |
| plastid_rada_mut_2 | 68615282 | 871.7 |
| plastid_rada_mut_3 | 45985626 | 1068.7 |
| plastid_rada_wild_1 | 53480887 | 234.2 |
| plastid_rada_wild_2 | 46684396 | 954.5 |
| plastid_rada_wild_3 | 46190084 | 776.4 |
| plastid_reca1_mut_1 | 66804365 | 543.7 |
| plastid_reca1_mut_2 | 38396319 | 594.8 |
| plastid_reca1_mut_3 | 30645358 | 299.3 |
| plastid_reca1_wild_1 | 37377457 | 598.2 |
| plastid_reca1_wild_2 | 32420159 | 543.5 |
| plastid_reca1_wild_3 | 33351491 | 331.1 |

**Table S3.** Results from Kruskal-Wallis test comparing SNV frequencies among genomic regions in WT and *msh1* mutant data from Wu *et al.,* (2020)

| Sample | Kruskal-Wallis chi-squared value | p-value |
| --- | --- | --- |
| *msh1* mitochondria | 6.03 | 0.19 |
| *msh1* plastid | 5.47 | 0.24 |
| WT mitochondria | 6.66 | 0.15 |
| WT plastid | 11.35 | 0.02 |

**Table S4.**
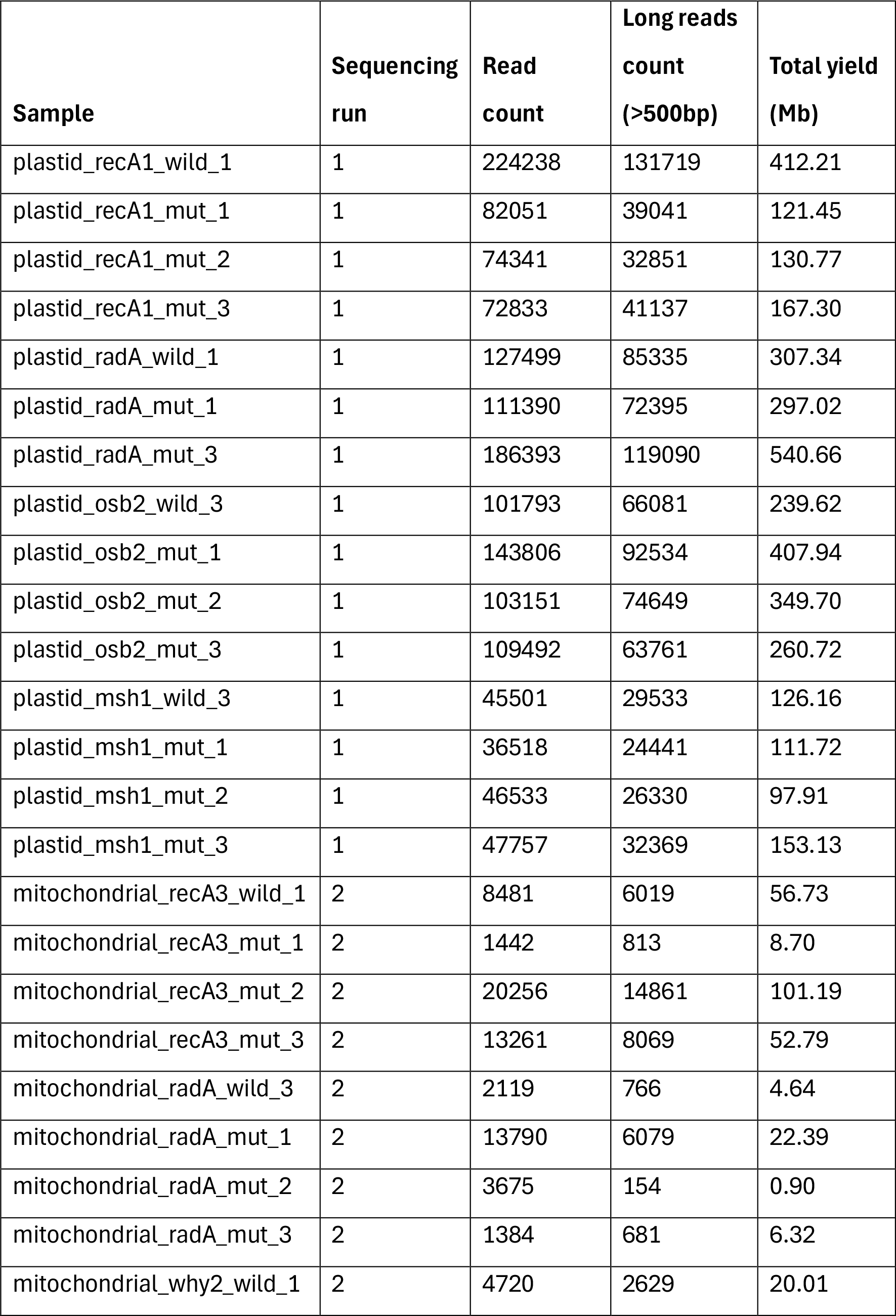
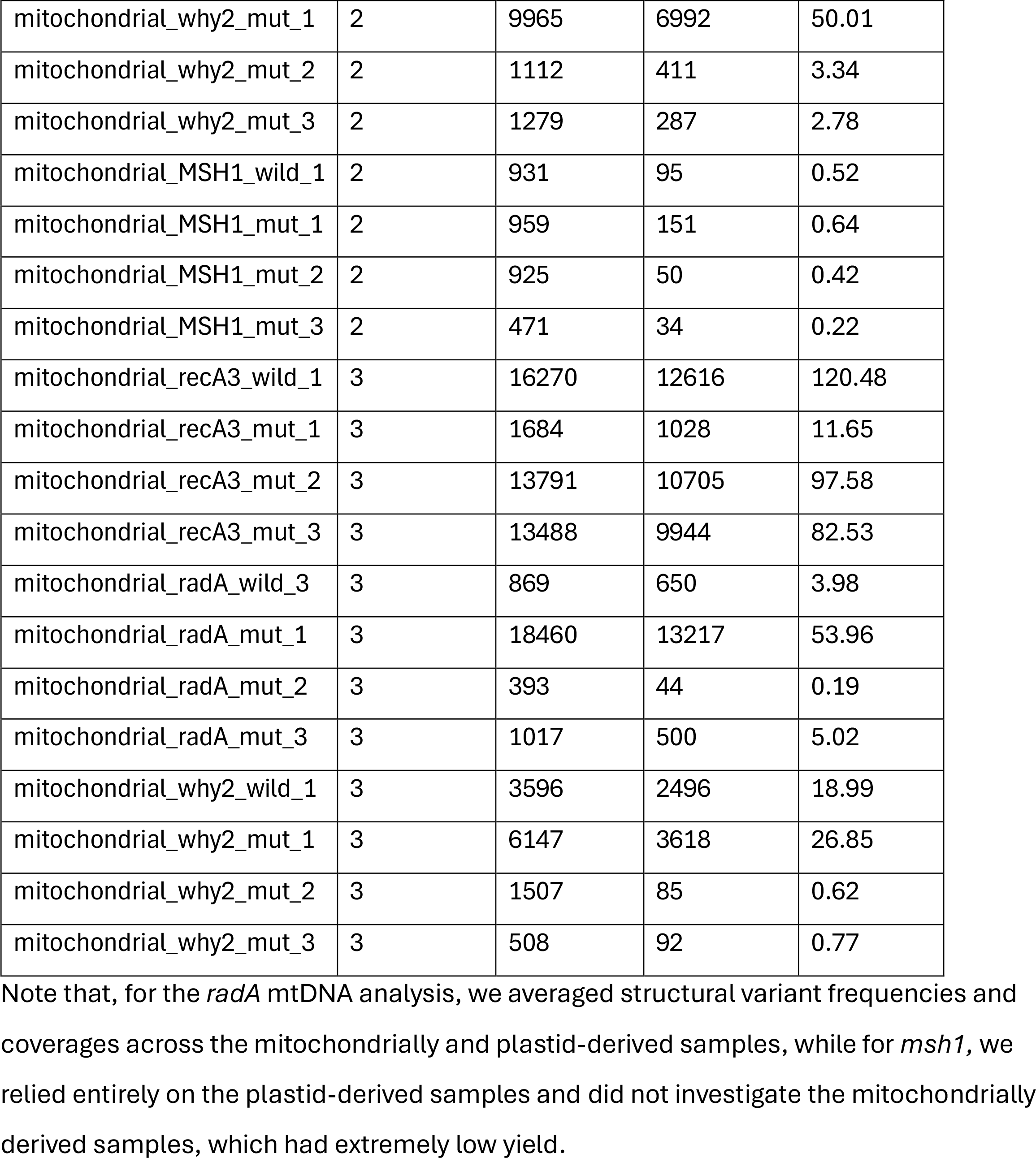
Oxford Nanopore sequencing yields for each of the three runs Note that, for the *radA* mtDNA analysis, we averaged structural variant frequencies and coverages across the mitochondrially and plastid-derived samples, while for *msh1,* we relied entirely on the plastid-derived samples and did not investigate the mitochondrially derived samples, which had extremely low yield.

**Table S5.** Sequencing depth per bp (calculated with bedtools depth) of samples in Fig 8.

| Sample | Sequencing protocol | Mutant (total cov/bp) | WT (total cov/bp) |
| --- | --- | --- | --- |
| radA_mito | nanopore | 222.9 | 101.6 |
| recA3_mito | nanopore | 736.4 | 372.1 |
| why2_mito | nanopore | 170.9 | 82.5 |
| msh1_mito | nanopore | 161.9 | 36.9 |
| radA_mito | duplex | 1600.2 | 1377.4 |
| recA3_mito | duplex | 941.4 | 647.8 |
| why2_mito | duplex | 1058.7 | 1235.2 |
| msh1_mito | duplex | 1020.9 | 1209.6 |
| msh1_plastid | nanopore | 1650.1 | 762.0 |
| osb2_plastid | nanopore | 6862.2 | 1663.7 |
| radA_plastid | nanopore | 5514.3 | 1668.5 |
| recA1_plastid | nanopore | 2422.6 | 2603.1 |
| msh1_plastid | duplex | 1754.4 | 1645.3 |
| osb2_plastid | duplex | 3244.9 | 3009.4 |
| radA_plastid | duplex | 3444.6 | 2205.6 |
| recA1_plastid | duplex | 1606.6 | 1650.2 |

## APPENDIX FOR SUPPLEMENTARY FILES

**FileS1_mutation_counts:** Coverages, mutation counts, and variant frequencies from the Duplex Sequencing analysis of data generated in this study and in Wu *et al.,* 2020.

**FileS2_repeat_recomb_freq_mito:** Counts of recombined reads and total repeat spanning reads used to calculate repeat specific recombination frequencies. We focused our mitochondrial analysis on repeats which has at least 10 recombined reads (across all replicates).

**FileS3_repeat_recomb_freq_plastid:** Counts of recombined reads and total repeat spanning reads used to calculate repeat specific recombination frequencies. We focused our plastid analysis on repeats which has at least 3 recombined reads (across all replicates).

